# Increased demand for NAD+ relative to ATP drives aerobic glycolysis

**DOI:** 10.1101/2020.06.08.140558

**Authors:** Alba Luengo, Zhaoqi Li, Dan Y. Gui, Lucas B. Sullivan, Maria Zagorulya, Brian T. Do, Raphael Ferreira, Adi Naamati, Ahmed Ali, Caroline A. Lewis, Craig J. Thomas, Stefani Spranger, Nicholas J. Matheson, Matthew G. Vander Heiden

## Abstract

Aerobic glycolysis, or preferential fermentation of glucose-derived pyruvate to lactate despite available oxygen, is a hallmark of proliferative metabolism that is observed across many organisms and conditions. To better understand why aerobic glycolysis is associated with cell proliferation, we examined the metabolic consequence of activating the pyruvate dehydrogenase complex (PDH) to increase mitochondrial pyruvate oxidation at the expense of fermentation. We find that increasing PDH activity impairs cell proliferation by reducing the nicotinamide adenine dinucleotide cofactor ratio (NAD+/NADH). This change in NAD+/NADH ratio is caused by an increase in mitochondrial membrane potential that impairs mitochondrial electron transport and NAD+ regeneration. Uncoupling mitochondrial respiration from ATP synthesis or increasing ATP hydrolysis restores NAD+/NADH homeostasis and proliferation even when glucose oxidation is increased. These data suggest that when the demand for NAD+ to support oxidation reactions exceeds the demand for ATP consumption in cells, NAD+ regeneration by mitochondrial respiration becomes constrained, promoting fermentation despite available oxygen. This argues that cells engage in aerobic glycolysis when the cellular demand for NAD+ is in excess of the cellular demand for ATP.

## Introduction

Cell growth and division impose increased energetic and biosynthetic demands, and thus proliferating cells exhibit a distinct metabolism relative to non-proliferating cells. Under aerobic conditions, most cells reduce oxygen to water via respiration to support the oxidation reactions used to derive energy from nutrients. When oxygen is limiting, cells instead ferment carbohydrates to generate a waste product such as lactate or ethanol as an alternative, less carbon efficient, way to derive energy from nutrients. However, some cells, including many rapidly proliferating cells, exhibit high rates of fermentation even when oxygen is abundant, a metabolic phenotype referred to as aerobic glycolysis. Aerobic glycolysis, also known as the Warburg effect in cancer cells, has been long associated with tumors (Koppenol, Bounds, & Dang, 2011; Otto Warburg, 1924; O. Warburg, 1956), but this phenotype is not unique to cancer. Aerobic glycolysis is exhibited by some proliferating microorganisms, including species of yeast and bacteria, and many proliferating non-transformed mammalian cells, including lymphocytes and fibroblasts (K. Brand, Leibold, Luppa, Schoerner, & Schulz, 1986; Hume, Radik, Ferber, & Weidemann, 1978; Lemoigne, Aubert, & Millet, 1954; Munyon & Merchant, 1959; Wang, Marquardt, & Foker, 1976). Select non-proliferative cells with high anabolic demands, such as pigmented epithelial cells of the mammalian retina, also engage in aerobic glycolysis (Chinchore, Begaj, Wu, Drokhlyansky, & Cepko, 2017; Krebs, 1927). Despite this being a phenotype found in many different cells and organisms, what drives aerobic glycolysis and why it is associated with proliferation has never been fully explained (Liberti & Locasale, 2016).

Aerobic glycolysis produces less ATP per mole of glucose than complete glucose oxidation to CO_2_, raising the question of why some cells engage in a metabolic program that is less efficient with respect to ATP generation (Koppenol et al., 2011; Liberti & Locasale, 2016; Vander Heiden, Cantley, & Thompson, 2009). This paradox is particularly apparent in cancer and led to the hypothesis that tumors must have defects in mitochondrial respiration (O. Warburg, 1956). Others have suggested that this phenotype is caused by tumor-associated hypoxia (Gatenby & Gillies, 2004); however, aerobic glycolysis is a feature of many cells without a precedent oxygen limitation (Vander Heiden et al., 2009). Tumors also retain functional mitochondria (Koppenol et al., 2011; Weinhouse, 1956) and require mitochondrial respiration for growth, progression, and metastasis (LeBleu et al., 2014; Tan et al., 2015; Viale et al., 2014; Weinberg et al., 2010). Collectively, these findings argue against mitochondrial damage or oxygen limitation as the primary driver of this phenotype. Furthermore, it is a common misconception that aerobic glycolysis involves suppression of oxidative phosphorylation (Yao et al., 2019). Aerobic glycolysis is best characterized by increased glucose uptake and increased fermentation with continued respiration, resulting in a shift in metabolism where more glucose is fermented relative to that which is oxidized.

Several other models have been put forth to explain why aerobic glycolysis is observed in proliferating cells. One proposal is that increased flux through glycolysis can shunt important biosynthetic precursors into anabolic reactions that branch from this pathway, contributing to production of nucleosides, lipids, and/or proteins (Boroughs & DeBerardinis, 2015; Cairns, Harris, & Mak, 2011; Hume & Weidemann, 1979; Levine & Puzio-Kuter, 2010; Vander Heiden et al., 2009). Though it is an attractive idea that inefficient ATP production is a tradeoff for supporting anabolic reactions, glycolytic intermediates are not necessarily elevated in proliferating cells (Lunt et al., 2015; Williamson et al., 1970) and many proliferating cells excrete the majority of glucose carbons as lactate. In fact, amino acids, rather than glucose, accounts for the majority of new carbon biomass of proliferating cells (Hosios et al., 2016). Another proposed benefit of the Warburg effect is increased ATP production, as many have argued that ATP can be generated with faster kinetics by aerobic glycolysis than by oxidative phosphorylation (Pfeiffer, Schuster, & Bonhoeffer, 2001). It has also been suggested that the energetic cost of synthesizing glycolytic enzymes is less than the cost of synthesizing components needed for respiration, such that ATP production by glycolysis confers a fitness advantage for cells (Basan et al., 2015). Nevertheless, why aerobic glycolysis is engaged in some, but not all, rapidly proliferating cells is not fully explained by existing models.

The end product of glycolysis is pyruvate, which can be fermented to lactate or further oxidized using a series of reactions that depend on mitochondrial respiration, in which the electrons released by glucose oxidation are disposed of via the reduction of oxygen to water. The first step in pyruvate oxidation is catalyzed by the pyruvate dehydrogenase complex (PDH), which converts pyruvate to acetyl-CoA in a reaction that is irreversible under physiological conditions. PDH activity is suppressed by a low NAD+/NADH ratio (Pettit, Pelley, & Reed, 1975) as well as by pyruvate dehydrogenase kinases (PDK) and pyruvate dehydrogenase phosphatases (PDP), which modulate PDH by inhibitory phosphorylation (Kolobova, Tuganova, Boulatnikov, & Popov, 2001; Korotchkina & Patel, 2001). Thus, PDH is a critical regulatory point for determining the extent to which cells engage in aerobic glycolysis (Grassian, Metallo, Coloff, Stephanopoulos, & Brugge, 2011; Kim, Tchernyshyov, Semenza, & Dang, 2006; Papandreou, Cairns, Fontana, Lim, & Denko, 2006). Activation of PDH either via inhibition of PDK activity or activation of PDP activity suppresses aerobic glycolysis, and can slow cancer cell proliferation and tumor growth (Hitosugi et al., 2011; Kaplon et al., 2013; McFate et al., 2008).

To better understand how aerobic glycolysis supports cell proliferation, we studied the consequence of suppressing fermentation in cells by increasing the activity of PDH. We find that promoting pyruvate oxidation impairs cell proliferation by limiting NAD+ availability for oxidation reactions, as interventions that regenerate NAD+ restore proliferation despite PDH activation. We further find that the reduced NAD+/NADH ratio in these cells results from an increase in mitochondrial membrane potential that impedes mitochondrial electron transport and NAD+ regeneration. Uncoupling electron transport from mitochondrial ATP synthesis relieves the increased mitochondrial membrane potential and increases NAD+ regeneration via respiration. Furthermore, increasing cellular ATP consumption rescues proliferation when PDH is activated, suggesting that ADP availability for coupled mitochondrial respiration can be an endogenous constraint of mitochondrial NAD+ regeneration. Lastly, endowing cells with alternative means of NAD+ regeneration suppresses aerobic glycolysis in both mammalian and yeast cells, without affecting proliferation rate. These data argue that cells engage in aerobic glycolysis when the NAD+ demand for oxidation reactions exceeds the demand for ATP and creates a situation where mitochondrial respiration is insufficient to support NAD+ regeneration.

## Results

### PDK inhibition activates PDH and suppresses aerobic glycolysis and cell proliferation

To better understand why proliferating cells engage in aerobic glycolysis, we sought to suppress this phenotype by increasing glucose oxidation relative to fermentation. Flux through PDH, the first committed step for mitochondrial glucose oxidation, is negatively regulated by PDK (Figure 1A), and PDK inhibition has been shown to suppress aerobic glycolysis in various contexts (Hitosugi et al., 2011; Kaplon et al., 2013; McFate et al., 2008; Michelakis et al., 2010). We utilized AZD7545, a potent and selective inhibitor of PDK1, PDK2, and PDK3 (Kato, Li, Chuang, & Chuang, 2007; Morrell et al., 2003), to promote pyruvate oxidation (Figure 1A). Exposing various cancer cells to AZD7545 decreased inhibitory phosphorylation of the E1α subunit of PDH (Figure 1B), confirming functional PDK inhibition by this small molecule. We next assessed PDH activity in cells with and without PDK inhibition by measuring incorporation of carbons from ^13^C-labeled glucose into citrate. We observed kinetics consistent with elevated PDH activity in cells treated with AZD7545 at multiple concentrations (Figure 1C, figure supplement 1A). Oxygen consumption rate (OCR) was also increased upon AZD7545 treatment in some, but not all, cells examined (Figure 1D, figure supplement 1B). However, AZD7545 treatment decreased the rate of lactate excretion per glucose molecule consumed in all tested cells (Figure 1E), confirming that PDK inhibition increases PDH activity, promoting pyruvate oxidation at the expense of fermentation, which represents a shift away from aerobic glycolysis.

**Figure 1.**
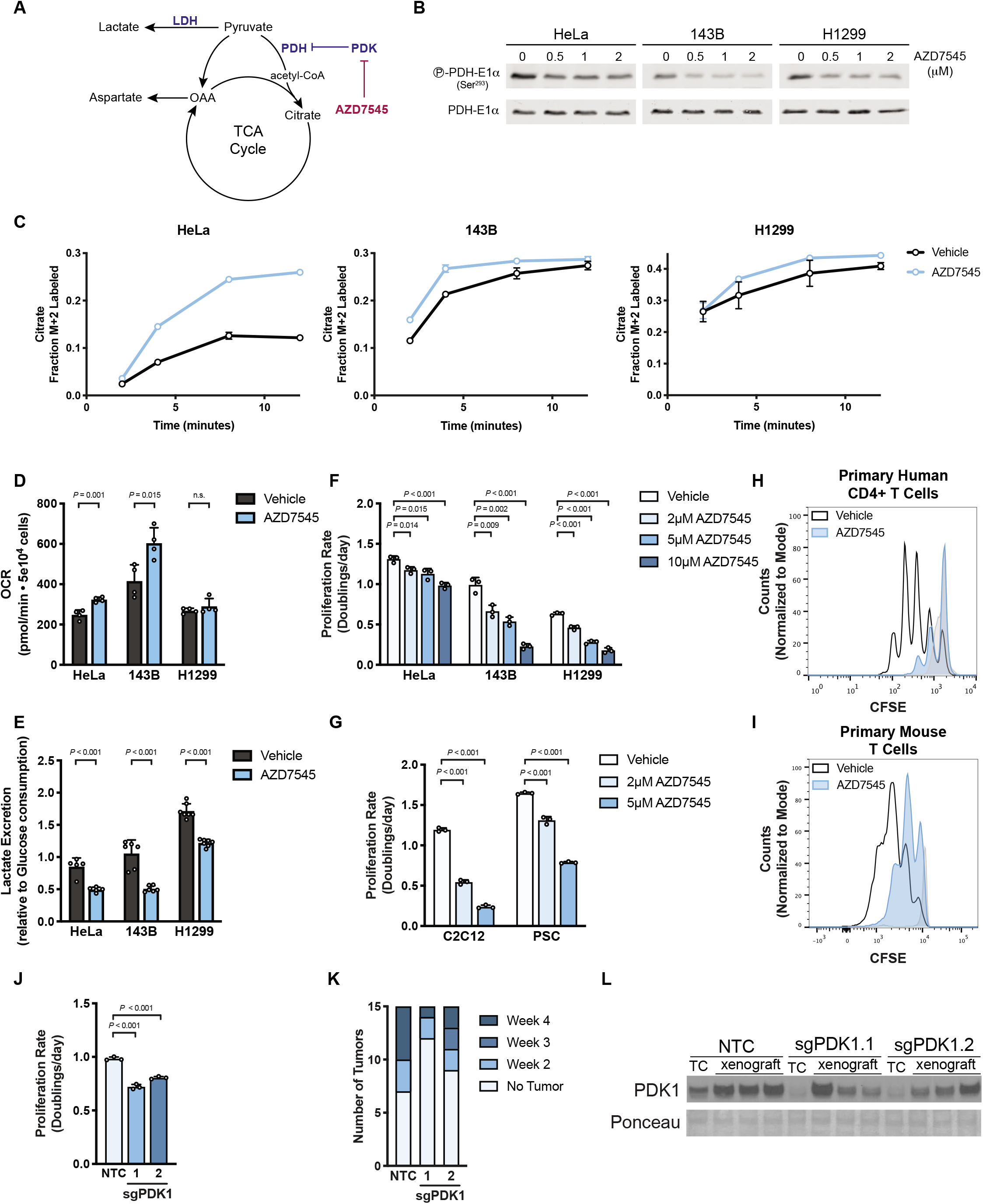
Activation of PDH suppresses aerobic glycolysis and proliferation. (A) Schematic illustrating the relationship between pyruvate dehydrogenase kinase (PDK) activity and pyruvate fate. Pyruvate has several fates in cells, including metabolism to lactate by lactate dehydrogenase (LDH) or oxidation by pyruvate dehydrogenase (PDH) to acetyl-CoA for entry into the tricarboxylic acid (TCA) cycle. PDH is under negative regulation of PDK enzymes, which can be inhibited by the compound AZD7545. (B) Western blot to assess S293 phosphorylation of the pyruvate dehydrogenase E1-alpha (P-PDH-E1α, Ser^293^) enzyme subunit in HeLa, 143B, and H1299 cells treated with vehicle or AZD7545 for 2 hours. Total PDH-E1α expression was also assessed. (C) Kinetic labeling of citrate from ^13^C-labeled glucose to assess PDH flux with and without PDK inhibition by AZD7545. HeLa, 143B and H1299 cells were incubated for 5 hours in media containing 5 mM unlabeled glucose with vehicle or 0.5 μM AZD7545, after which 20 mM [U-^13^C_6_]glucose was added. The fraction of M+2 citrate was measured by LCMS after the addition of ^13^C-labeled glucose (n = 3 per time point). (D) Oxygen consumption rate (OCR) of cells that had been treated with vehicle or 0.5 μM AZD7545 for 5 hours (n = 4). (E) Lactate excretion into culture media normalized to glucose consumption of cells treated with vehicle or 0.25 μM AZD7545 for 48 hours (n = 5). (F) Proliferation rate of 143B, H1299, and HeLa cells treated with vehicle or the indicated concentration of AZD7545 (n = 3). (G) Proliferation rate of C2C12 myoblasts or mouse pancreatic stellate cells (PSC) cultured in media containing vehicle or the indicated concentration of AZD7545 (n = 3). (H) Proliferation of primary human CD4+ T cells in the presence of vehicle or 5 μM AZD7545. Human CD4+ T cells were isolated and stained with CFSE prior to stimulation with anti-CD3/CD28 dynabeads, and CFSE fluorescence was assessed by flow cytometry after 4 days. Representative data are shown from 3 biological replicates of primary human CD4+ T cells collected from different donors and analyzed as independent experiments. Data from CFSE-stained, unstimulated cells (light grey) are also shown as a control for fluorescence of cells that did not proliferate. (I) Proliferation of primary mouse T cells in the presence of vehicle or 5 μM AZD7545. Mouse T cells were isolated and stained with CFSE prior to stimulation with anti-CD3/CD28 antibodies, and CFSE fluorescence was assessed by flow cytometry after 2 days. Stained, unstimulated cells (light grey) are shown as a control for fluorescence of cells that did not proliferate. (J) Proliferation rate of 143B cells in which CRISPR interference was used to transcriptionally repress PDK1 expression. Cells were transduced with sgRNA targeting PDK1 (sgPDK1; two independently targeted lines) or a non-targeting control (NTC) as indicated (n = 3). (K) Histogram indicating the number of weeks at which the cell lines described in (J) formed xenograft tumors larger than 50 mm^3^ in nude mice (n = 15). (L) Western blot analysis to assess PDK1 expression in the cells shown in panel (J) when cultured in vitro (TC) or after being isolated from the xenografts described in (K) after 34 days. Values shown in panels C, D, E, F, G, and J denote the mean ± SD and the indicated *P* values were calculated by unpaired, two-tailed Student’s t-test (n.s. = not significant).

We next tested the effects of AZD7545 on cell proliferation and found that suppression of aerobic glycolysis decreased proliferation of both cancer cells (Figure 1F) and non-transformed C2C12 myoblast cells and primary murine pancreatic stellate cells (PSCs) (Figure 1G). AZD7545 also suppressed proliferation of activated primary human CD4+ T cells and primary mouse T cells (Figure 1H,I; figure supplement 1C,D) while having minimal effect on the expression of T cell activation markers (figure supplement 1E). These data demonstrate that PDH activation impairs proliferation in both cancer and non-cancer settings.

To confirm that AZD7545 inhibits cell proliferation as a result of PDK inhibition rather than another effect of this compound, we disrupted PDK1 expression in 143B cells using CRISPR interference (CRISPRi) (figure supplement 1F) and found that genetic suppression of PDK1 slowed cell proliferation (Figure 1J). Furthermore, high PDH activity is selected against for tumor growth in mice, as 143B cells with PDK1 knockdown displayed impaired tumor formation relative to controls (Figure 1K). Of note, when PDK1 knockdown cells form tumors, some of these tumors grow at a similar rate as tumors derived from control cells (figure supplement 1G); however, tumors derived from PDK1 knockdown cells regained PDK1 expression (Figure 1L). Furthermore, control and PDK1 knockdown tumors expressed higher levels of PDK1 than the cells from which they were derived (Figure 1L). Taken together, these data are consistent with numerous studies showing enhanced PDH activation specifically (Kaplon et al., 2013; McFate et al., 2008; Michelakis et al., 2010), and suppressing aerobic glycolysis more generally (Fantin, St-Pierre, & Leder, 2006; Le et al., 2010; Xie et al., 2014), can slow cell proliferation and tumor growth.

### PDH activation decreases proliferation by decreasing the NAD+/NADH ratio in cells

A better understanding of how increased pyruvate oxidation suppresses cell proliferation could provide insight into why proliferating cells exhibit aerobic glycolysis. One potential explanation is that altering the ratio of glucose oxidation to lactate fermentation can affect the oxidative capacity of cells. The NAD+/NADH ratio is critical for many metabolic reactions, including those involved in central carbon metabolism, nucleotide synthesis, lipid metabolism, and amino acid metabolism (Hosios & Vander Heiden, 2018), and NAD+ regeneration can be limiting for cell proliferation and tumor growth (Birsoy et al., 2015; Gui et al., 2016; Sullivan et al., 2015; Titov et al., 2016). The reaction catalyzed by PDH consumes NAD+ to produce NADH, and shunting pyruvate away from lactate production prevents NAD+ regeneration by lactate dehydrogenase (LDH) (Figure 2A). Thus, PDK inhibition is expected to favor a more reduced cellular NAD+/NADH ratio, and indeed AZD7545 treatment lowers the NAD+/NADH ratio in cells (Figure 2B, figure supplement 2A).

**Figure 2.**
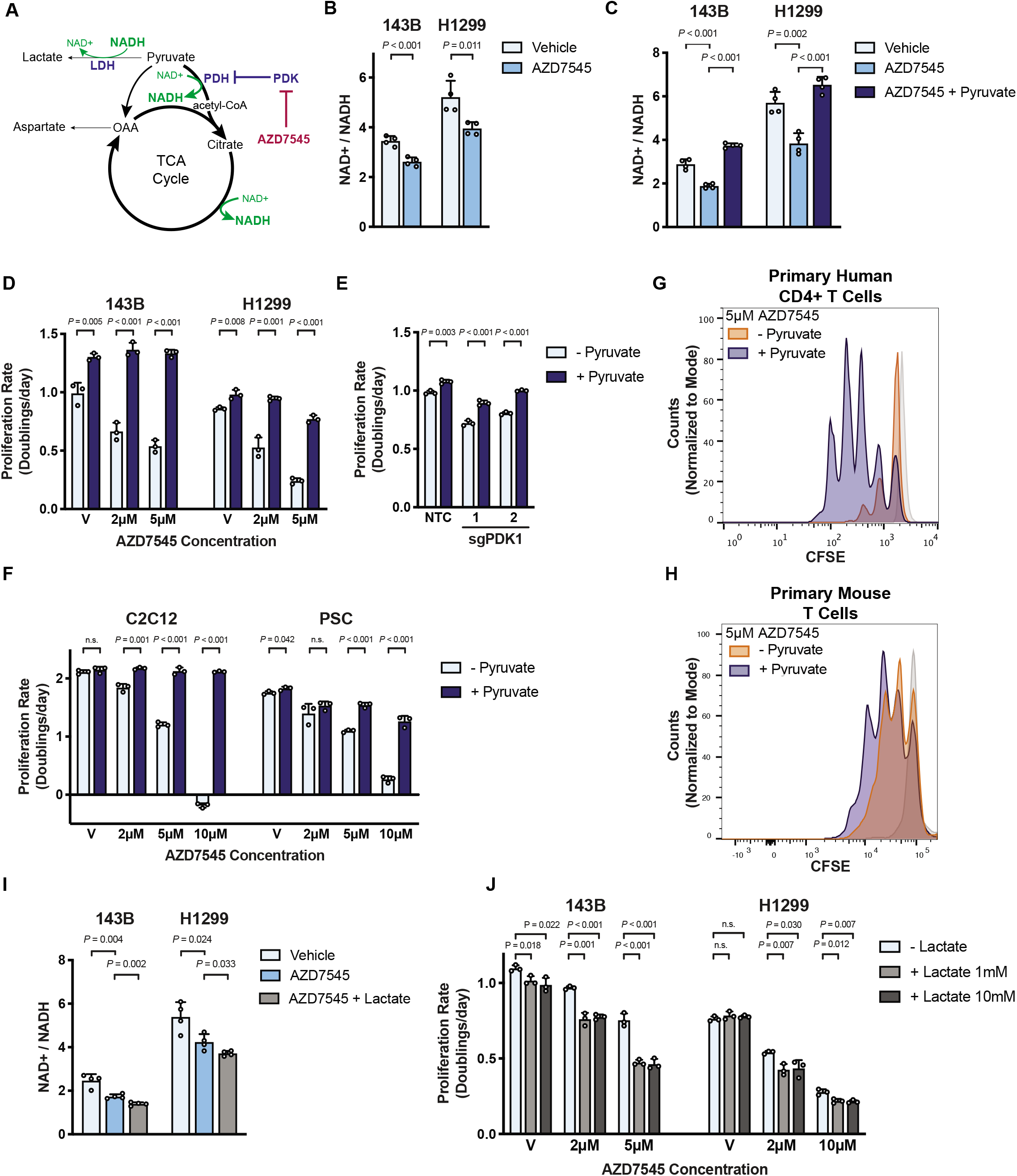
PDK inhibition slows cell proliferation by reducing the NAD+/NADH ratio. (A) Schematic depicting the redox consequences of PDK inhibition. PDK inhibition by AZD7545 decreases the NAD+/NADH ratio by promoting flux through NAD+ consuming pathways, including PDH and the TCA cycle, while limiting pyruvate conversion to lactate by LDH. (B) NAD+/NADH ratio of 143B and H1299 cells cultured in the presence of vehicle or 5 μM AZD7545 for 5 hours (n = 4). (C) NAD+/NADH ratio of 143B and H1299 cells treated with vehicle, 5 μM AZD7545, or 5 μM AZD7545 with 1 mM pyruvate (n = 4). (D) Proliferation rate of 143B and H1299 cells cultured in vehicle (V) or the indicated concentration of AZD7545 in the presence (dark blue) or absence (light blue) of 1 mM pyruvate (n = 3). (E) Proliferation rate of 143B cells in which CRISPR interference (CRISPRi) was used to transcriptionally repress PDK1. Cells were transduced with sgRNA targeting PDK1 or a nontargeting control (NTC) and grown in the presence (dark blue) or absence (light blue) of 1 mM pyruvate (n = 3). (F) Proliferation rate of C2C12 myoblasts and pancreatic stellate cells (PSC) grown in media containing vehicle (V) or AZD7545 supplemented with (dark blue) or without (light blue) 1 mM pyruvate (n = 3). (G) Proliferation of primary human CD4+ T cells treated with 5 μM AZD7545 in the presence or absence of 1 mM pyruvate. Human CD4+ T cells were isolated and stained with CFSE prior to stimulation with CD3/CD28 dynabeads, and CFSE fluorescence was assessed by flow cytometry after 4 days. Representative data are shown from 3 biological replicates of primary human CD4+ T cells collected from different donors and analyzed as independent experiments. Stained, unstimulated cells (light grey) are shown as a control for fluorescence of cells that did not proliferate. (H) Proliferation of primary mouse T cells treated with 5 μM AZD7545 in the presence or absence of 1 mM pyruvate. Mouse T cells were isolated and stained with CFSE prior to stimulation with anti-CD3/CD28 antibodies. CFSE fluorescence was assessed by flow cytometry after 2 days. Stained, unstimulated cells (light grey) are shown as a control for fluorescence of cells that did not proliferate. (I) NAD+/NADH ratio of 143B and H1299 cells cultured with vehicle, 5 μM AZD7545, or 5 μM AZD7545 with 10 mM lactate (n = 4). (J) Proliferation rate of 143B and H1299 cells treated with vehicle (V) or AZD7545 with or without 1 mM or 10 mM lactate as indicated (n = 3). Values shown in panels B, C, D, E, F, I and J denote the mean ± SD. *P* values shown were calculated by unpaired, two-tailed Student’s t-test unless otherwise indicated (n.s. = not significant).

We next questioned whether sensitivity to PDK inhibition was affected by altering the intracellular NAD+/NADH ratio. Pyruvate is rapidly reduced by LDH, and providing exogenous pyruvate increases the NAD+/NADH ratio (Birsoy et al., 2015; Gui et al., 2016; Sullivan et al., 2015). Pyruvate supplementation restored the NAD+/NADH ratio in cells treated with AZD7545 (Figure 2C) and supplying exogenous pyruvate enhanced baseline cancer cell proliferation and suppressed the anti-proliferative effects of AZD7545 as well as genetic suppression of PDK1 (Figure 2D,E; figure supplement 2B). Pyruvate also increased the proliferation rate of AZD7545-treated non-transformed C2C12 cells and mouse PSCs (Figure 2F), and of activated primary human CD4+ T cells (Figure 2G, figure supplement 2C,D) and primary mouse T cells (Figure 2H; figure supplement 2E,F). These data argue that providing exogenous pyruvate suppresses the anti-proliferative effect of PDH activation.

Lactate can serve as a nutrient for cells, including cancer cells (Faubert et al., 2017; Hui et al., 2017; Kennedy et al., 2013; Sonveaux et al., 2008). Lactate metabolism first requires conversion to pyruvate where it serves as an electron donor for the LDH reaction and converts NAD+ to NADH (Figure 2A). Thus, supplying exogenous lactate further decreased the NAD+/NADH ratio of cells treated with AZD7545 (Figure 2I, figure supplement 2G). Exogenous lactate also suppressed cell proliferation and exacerbated the effect of AZD7545 treatment in 143B and H1299 cells (Figure 2J), but not HeLa cells (figure supplement 2H). Titrating extracellular lactate relative to extracellular pyruvate has been used to titrate the NAD+/NADH ratio in cells (Hung, Albeck, Tantama, & Yellen, 2011), and we found that lower extracellular lactate to pyruvate ratios suppressed the ability of AZD7545 to impair proliferation (figure supplement 2I). These data support depletion of NAD+ as an explanation for why increasing pyruvate oxidation slows proliferation.

To more directly test whether changes in the NAD+/NADH ratio mediate the anti-proliferative effects of PDK inhibition, we tested whether orthogonal pathways that regenerate NAD+, but are not involved in glucose, pyruvate, or lactate metabolism, affect sensitivity to AZD7545. Duroquinone permits NAD+ regeneration via the quinone reductase NQO1 (Merker, Audi, Bongard, Lindemer, & Krenz, 2006) (Figure 3A) and duroquinone can rescue cell proliferation in conditions where NAD+ is limiting (Gui et al., 2016). We found that duroquinone suppresses AZD7545 sensitivity (Figure 3B, figure supplement 3A), consistent with a decreased NAD+/NADH ratio slowing proliferation of PDK-inhibited cells.

**Figure 3.**
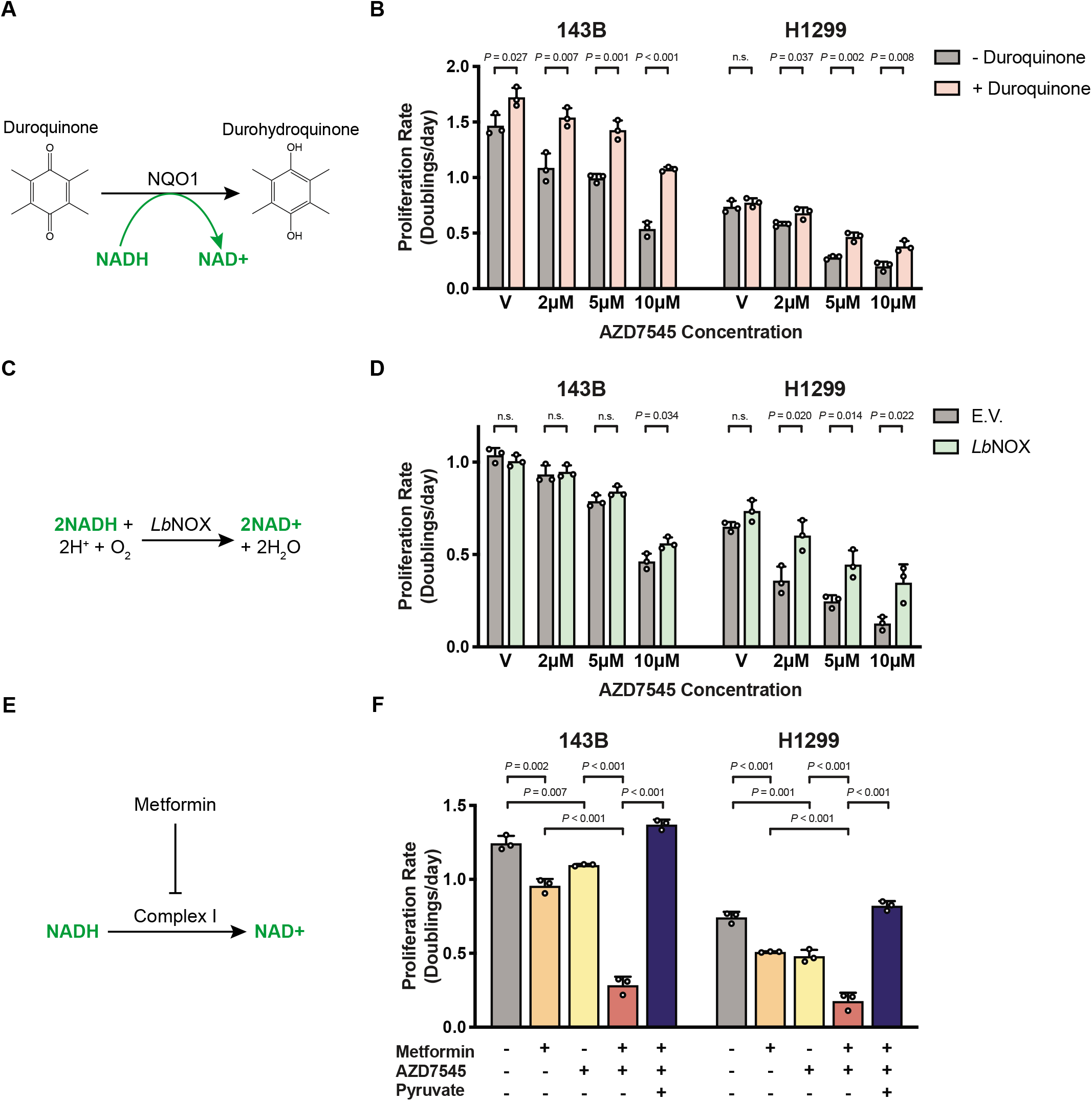
Interventions that alter NAD+ availability can modulate the antiproliferative effects of PDK inhibition. (A) Schematic illustrating how duroquinone supplementation increases NAD+ regeneration via the enzyme NAD(P)H dehydrogenase, quinone 1 (NQO1), which reduces duroquinone to durohydroquinone using NADH as a cofactor. (B) Proliferation rate of 143B and H1299 cells treated with vehicle (V) or AZD7545 in the absence or presence of duroquinone. The duroquinone concentration used was 20 μM for 143B and 100 μM for H1299 cells (n = 3). (C) Schematic illustrating the reaction catalyzed by the NADH oxidase from *Lactobacillus brevis* (*Lb*NOX). (D) Proliferation rate of 143B and H1299 cells transduced with empty vector (E.V.) or an *Lb*NOX expression vector and treated with vehicle (V) or AZD7545. Doxycycline (500 ng/mL) was included in all conditions to induce *Lb*NOX expression (n = 3). (E) Schematic illustrating the redox consequences of metformin treatment. (F) Proliferation rate of 143B and H1299 cells treated with 500 μM metformin, AZD7545 (5 μM for 143B, 3 μM for H1299), and 1 mM pyruvate as indicated (n = 3). Values shown in panels B, D, and F denote the mean ± SD. *P* values were calculated by unpaired, two-tailed Student’s t-test (n.s. = not significant).

Another orthogonal method for increasing the cell NAD+/NADH ratio is expressing NADH oxidase from *Lactobacillus brevis* (*Lb*NOX) (Titov et al., 2016) (Figure 3C). Thus, we engineered cells to express either *Lb*NOX or empty vector (E.V.) as another way to assess the effect of increasing NAD+ regeneration in cells (figure supplement 3B). *Lb*NOX expression conferred resistance to the anti-proliferative effects of AZD7545 (Figure 3D, figure supplement 3C), further demonstrating that increasing NAD+ regeneration can permit rapid proliferation despite increased pyruvate oxidation.

### Alterations in NAD+/NADH account for how PDH activation impacts cell metabolism

NAD+ is necessary to support oxidation reactions in cells, and thus changes in NAD+/NADH ratio impacts many metabolic pathways including those important for cell growth and proliferation (Hosios & Vander Heiden, 2018). For example, the NAD+/NADH ratio can affect de novo aspartate synthesis (Birsoy et al., 2015; Sullivan et al., 2015). Aspartate is essential to make proteins as well as purine and pyrimidine nucleotides, and acquiring aspartate can be limiting for tumor growth (Garcia-Bermudez et al., 2018; Rabinovich et al., 2015; Sullivan et al., 2018). Therefore, to determine whether PDK inhibition affects aspartate availability as one way to impair proliferation, we assessed aspartate levels in cells cultured in the presence of vehicle or AZD7545. Consistent with AZD7545 treatment reducing the cell NAD+/NADH ratio, PDK inhibition decreased intracellular aspartate (figure supplement 3D), and increasing the NAD+/NADH ratio with exogenous pyruvate, duroquinone, or *LbNOX* expression restored aspartate levels (figure supplement 3E-G). These data suggest that suppressing aerobic glycolysis limits the capacity of cells to carry out oxidation reactions that are important to produce biomass.

To further study how PDH activation affects metabolism, we performed untargeted metabolomics and observed that despite an increase in PDH activity (Figure 1C, figure supplement 1A), AZD7545 treatment decreased intracellular citrate levels and increased intracellular palmitate levels (figure supplement 3H,I). This could reflect a metabolic state where forcing cells to produce citrate from pyruvate will elicit a compensatory increase in de novo lipid synthesis, which consumes NADPH to make NADP+, potentially as an alternative means to regenerate oxidizing equivalents (Liu et al., 2020). We also observed that AZD7545 treatment affects the levels of many other intracellular metabolites, and that either duroquinone or expression of *Lb*NOX can suppress some of these changes (figure supplement 3J-M). Of note, duroquinone or *Lb*NOX expression restore many of the same metabolites in AZD7545-treated cells (figure supplement 3N). Because duroquinone and *Lb*NOX alter NAD+/NADH ratios via different mechanisms, the finding that both agents similarly restore alterations to the global metabolome caused by PDH activation argues that an effect on cell redox state is a major consequence of increased pyruvate oxidation and further supports the notion that a major metabolic consequence of suppressing aerobic glycolysis is NAD+ depletion.

### PDH activation increases dependency on mitochondrial complex I for NAD+ regeneration

Since PDK inhibition reduces the cellular NAD+/NADH ratio, interventions that limit NAD+ regeneration are predicted to potentiate the anti-proliferative effects of PDK inhibitors. To test this hypothesis, we assessed the sensitivity of cancer cells to the biguanide metformin, which limits NAD+ regeneration via inhibition of mitochondrial complex I (Gui et al., 2016; Wheaton et al., 2014)(Figure 3E). The combination of metformin and AZD7545 reduced cell proliferation more than either compound alone (Figure 3F, figure supplement 4A), and AZD7545 treatment decreased the IC_50_ of metformin by more than 30% in both A549 and HeLa cells (figure supplement 4B,C). The anti-proliferative effect of AZD7545 and metformin was completely abolished by pyruvate supplementation (Figure 3F, figure supplement 4A-C), further arguing that these drugs impair proliferation by promoting a more reduced NAD+/NADH ratio.

Electron acceptor availability can be limiting for tumor growth in some mouse models of cancer (Gui et al., 2016), suggesting that inhibiting NAD+ regeneration with the combination of AZD7545 and metformin may have a larger effect on tumor growth than either drug alone. We found that metformin inhibited tumor growth as previously reported (Gui et al., 2016; Schockel et al., 2015; Wheaton et al., 2014), and while AZD7545 had no effect alone, PDK inhibition appeared to improve the efficacy of metformin in this model (figure supplement 4D).

The finding that AZD7545 treatment did not decrease tumor growth as a single agent might be expected from the pharmacokinetics of this compound, as well as the fact that AZD7545 impairs, but does not prevent, cell proliferation. Three hours after dosing, we find levels of AZD7545 in tumors and in serum that can impair proliferation of cells in culture (figure supplement 4E,F), but the compound was not detected in plasma after 24 hours (figure supplement 4E). This argues that more frequent dosing of AZD7545 may be more effective to inhibit tumor growth as a single agent. Nevertheless, once a day dosing of AZD7545 was sufficient to improve the anti-tumor effects of metformin, supporting the notion that PDK inhibition increases dependency on mitochondrial complex I to regenerate NAD+ and support proliferation.

### Increased mitochondrial membrane potential downstream of pyruvate oxidation impairs NAD+ regeneration by the mitochondrial electron transport chain

The decreased NAD+/NADH ratio observed in PDK-inhibited cells suggests that NAD+ regeneration by mitochondrial respiration is insufficient to support maximal proliferation when pyruvate oxidation is increased. This result is unexpected, as these cells are cultured at atmospheric oxygen and exhibit an increased rate of basal oxygen consumption upon AZD7545 treatment (Figure 1D). Thus, we sought to better understand how NAD+ regeneration by mitochondrial respiration was impaired in PDK-inhibited cells, as this could explain why proliferating cells engage in aerobic glycolysis.

The oxidation-reduction reactions of the mitochondrial electron transport chain (ETC) are coupled to proton pumping from the mitochondrial matrix into the intermembrane space to generate an electrochemical gradient across the inner mitochondrial membrane and support ATP production via the FoF1-ATP synthase (Figure 4A). This process, collectively referred to as oxidative phosphorylation, occurs at near equilibrium, meaning that the rate of respiration is a function of both substrate and product availability (Brown, 1992). The shift toward a more reduced NAD+/NADH ratio suggests NADH is more readily available in these cells, but one possibility is that the availability of oxygen, another major substrate for oxidative phosphorylation, becomes limiting for electron transport. However, these cells were cultured at 21% oxygen, and mitochondrial respiration can function at oxygen levels as low as 0.5% (Chandel, Budinger, & Schumacker, 1996; Rumsey, Schlosser, Nuutinen, Robiolio, & Wilson, 1990). Furthermore, expression of *Lb*NOX, which requires oxygen as a substrate, suppressed the anti-proliferative effect of AZD7545 (Figure 3D), arguing against oxygen availability constraining NAD+ regeneration in PDK-inhibited cells.

**Figure 4.**
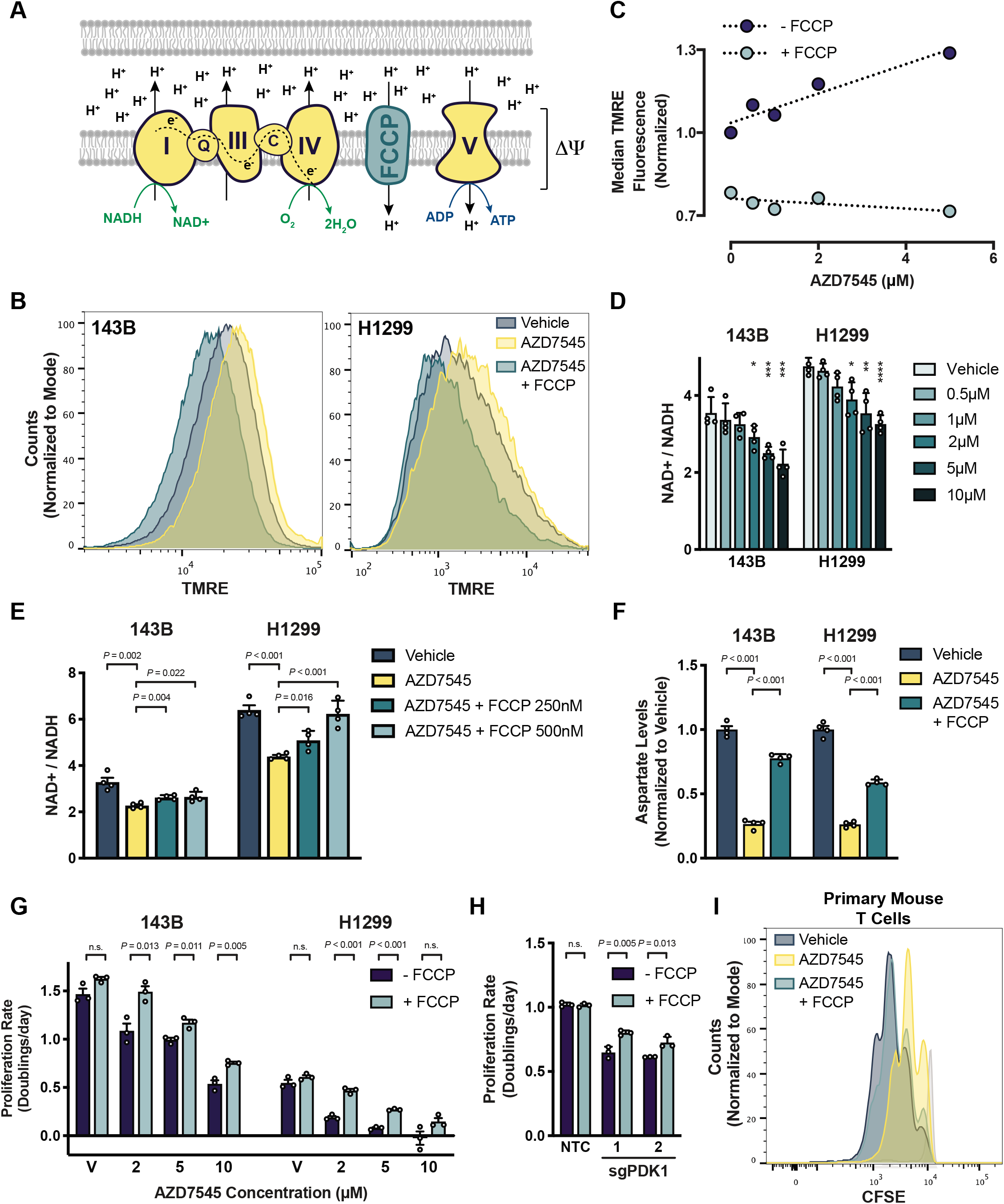
PDK inhibition induces mitochondria hyperpolarization and limits NAD+ regeneration by respiration. (A) Schematic illustrating the mitochondrial electron transport chain and how FCCP (trifluoromethoxy carbonylcyanide phenylhydrazone) uncouples electron transfer from NADH to O_2_ from ATP production by the F_o_F_1_-ATP synthase (Complex V). ΔΨ denotes the mitochondrial membrane potential. (B) Mitochondrial membrane potential, as reflected by TMRE (tetramethylrhodamine, ethyl ester) fluorescence, in 143B and H1299 cells treated with vehicle, 5 μM AZD7545, or 5 μM AZD7545 with 500 nM FCCP. (C) Mitochondrial membrane potential, as reflected by TMRE fluorescence, in H1299 cells that had been treated with the indicated concentration of AZD7545, with or without 500 nM FCCP. TMRE fluorescence of 50,000 cells was quantified using flow cytometry is normalized to the vehicle-treated condition without FCCP as shown. (D) NAD+/NADH ratio of 143B and H1299 cells cultured in vehicle or AZD7545 for 5 hours (n = 4). (E) NAD+/NADH ratio of 143B and H1299 cells treated with vehicle, 5 μM AZD7545, or 5 μM AZD7545 with the indicated concentration of FCCP for 5 hours (n = 4). (F) Aspartate levels in 143B and H1299 cells that were cultured with vehicle, 2 μM AZD7545, or 2 μM AZD7545 with 250 nM FCCP for 5 hours as measured by LCMS (n = 4). (G) Proliferation rate of 143B and H1299 cells treated with vehicle (V) or AZD7545 in the absence or presence of 500 nM FCCP as indicated (n = 3). (H) Proliferation rate of 143B cells in which CRISPR interference was used to transcriptionally repress PDK1. Cells were transduced with sgRNA targeting PDK1 (sgPDK1; two independently targeted lines) or a non-targeting control (NTC) and cultured in the presence or absence of 750 nM FCCP (n = 3). (I) Proliferation of primary mouse T cells in the presence of vehicle, 5 μM AZD7545, or 5 μM AZD7545 with 500 nM FCCP was assessed by CFSE dye dilution. Mouse T cells were isolated and stained with CFSE prior to stimulation with anti-CD3/CD28 antibodies and CFSE fluorescence was assessed by flow cytometry after 2 days. Stained, unstimulated cells (light grey) are shown as a control for fluorescence of cells that did not proliferate. Values in panel D, E, F, G, and H denote the mean ± SD. *P* values shown were calculated by unpaired, two-tailed Student’s t-test. *, **, ***, and **** denote *P* < 0.05, 0.01, 0.005, and 0.001 respectively (n.s. = not significant).

Another possibility is that an imbalance develops between the ability of the mitochondrial ETC to pump protons into the intermembrane space and the activity of processes that carry protons back across the membrane. This imbalance would be reflected in an increased mitochondrial membrane potential (ΔΨ)(Figure 4A). To test this possibility, we utilized the cationic dye tetramethylrhodamine ethyl ester (TMRE), which is taken up by mitochondria in proportion to ΔΨ (Perry, Norman, Barbieri, Brown, & Gelbard, 2011). We found that PDK inhibition increases TMRE accumulation in cancer cells (Figure 4B) and activated primary mouse T cells (figure supplement 5A), and that this effect is reversed when respiration is uncoupled from ATP production by the ionophore FCCP (trifluoromethoxy carbonylcyanide phenylhydrazone)(Figure 4B). FCCP allows protons to equilibrate across membranes, thus FCCP uncouples NAD+ regeneration by the ETC from ΔΨ generation and prevents use of ΔΨ to produce ATP (Figure 4A). These data are consistent with PDH activation increasing ΔΨ, and increased TMRE accumulation caused by AZD7545 treatment depends on the concentration of AZD7545 (Figure 4C) in a way that matches the dose-dependent reduction in the NAD+/NADH ratio observed upon PDK inhibition (Figure 4D, figure supplement 5B). Taken together, these data suggest that NAD+ regeneration by respiration is constrained by ΔΨ, as it becomes thermodynamically unfavorable for the mitochondrial ETC complexes to pump protons across a hyperpolarized mitochondrial membrane.

If high ΔΨ limits NAD+ regeneration by respiration, collapse of ΔΨ should restore NAD+/NADH homeostasis to cells with activated PDH. Indeed, FCCP exposure was sufficient to increase the NAD+/NADH ratio (Figure 4E), aspartate levels (Figure 4F, figure supplement 5C) and proliferation rate (Figure 4G, figure supplement 5D) of PDK-inhibited cells. FCCP treatment also reversed the proliferation defect observed in cancer cells where PDK1 expression is suppressed using CRISPRi (Figure 4H). Additionally, FCCP treatment was capable of rescuing the proliferation rates of AZD7545-treated primary mouse T cells (Figure 4I), but not of primary human T cells. These results are consistent with ΔΨ limiting mitochondrial NAD+ regeneration. In addition, the finding that FCCP increases respiration confirms that the mitochondrial ETC machinery is functional and that sufficient oxygen is available to support increased respiration in these cells. Taken together, these data argue that mitochondrial NAD+ regeneration becomes limited by an increase in ΔΨ when pyruvate oxidation is increased.

### NAD+ regeneration is limited by excess ATP when PDK is inhibited

A physiological function of ΔΨ is to support ATP synthesis, as conversion of ADP to ATP by the F_o_F_1_-ATP synthase is coupled to dissipation of ΔΨ (Figure 4A). Thus, an excess of cellular ATP that limits ADP availability to support oxidative phosphorylation may explain why mitochondrial NAD+ regeneration is impaired in cells with high pyruvate dehydrogenase activity. The observation that FCCP treatment promotes proliferation of PDK-inhibited cells, while uncoupling mitochondrial NAD+ regeneration from ATP synthesis, is consistent with this model. This model also predicts that increasing ATP consumption in cells should increase ADP availability to support increased respiration and reverse the metabolic and anti-proliferative effects of PDK inhibition.

Promoting cellular ATP consumption stimulates mitochondrial respiration by increasing the demand for ADP to ATP conversion (Bertholet et al., 2019; Brown, 1992). The toxin gramicidin D is a classic way to increase ATP consumption in cells by increasing cell membrane permeability to Na+ and K+ ions, driving increased ATP hydrolysis by Na+/K+-ATPase and thus increasing ATP-coupled mitochondrial respiration (Buttgereit & Brand, 1995; Nobes, Lakin-Thomas, & Brand, 1989; Vander Heiden, Chandel, Schumacker, & Thompson, 1999). Thus, we used gramicidin D to determine whether increased ATP consumption affects the metabolic and anti-proliferative effects of PDK inhibition. First, we confirmed that increasing cellular ATP consumption with gramicidin D reversed mitochondrial membrane hyperpolarization caused by PDK-inhibition (Figure 5A). Exposing PDK-inhibited cells to gramicidin D also resulted in a more oxidized NAD+/NADH ratio (figure supplement 5E). These data are consistent with gramicidin D causing an increase in mitochondrial ATP synthesis, which in turn promotes increased respiration and mitochondrial NAD+ regeneration. Of note, gramicidin D also renders both cancer cells and non-cancer cells more resistant to AZD7545 treatment (Figure 5B,C). Taken together, these data support a model where mitochondrial NAD+ regeneration via respiration is limited by excess cellular ATP when pyruvate oxidation is increased. These findings also suggest that cells may engage in aerobic glycolysis under conditions where the demand for NAD+ regeneration exceeds the rate of ATP consumption.

**Figure 5.**
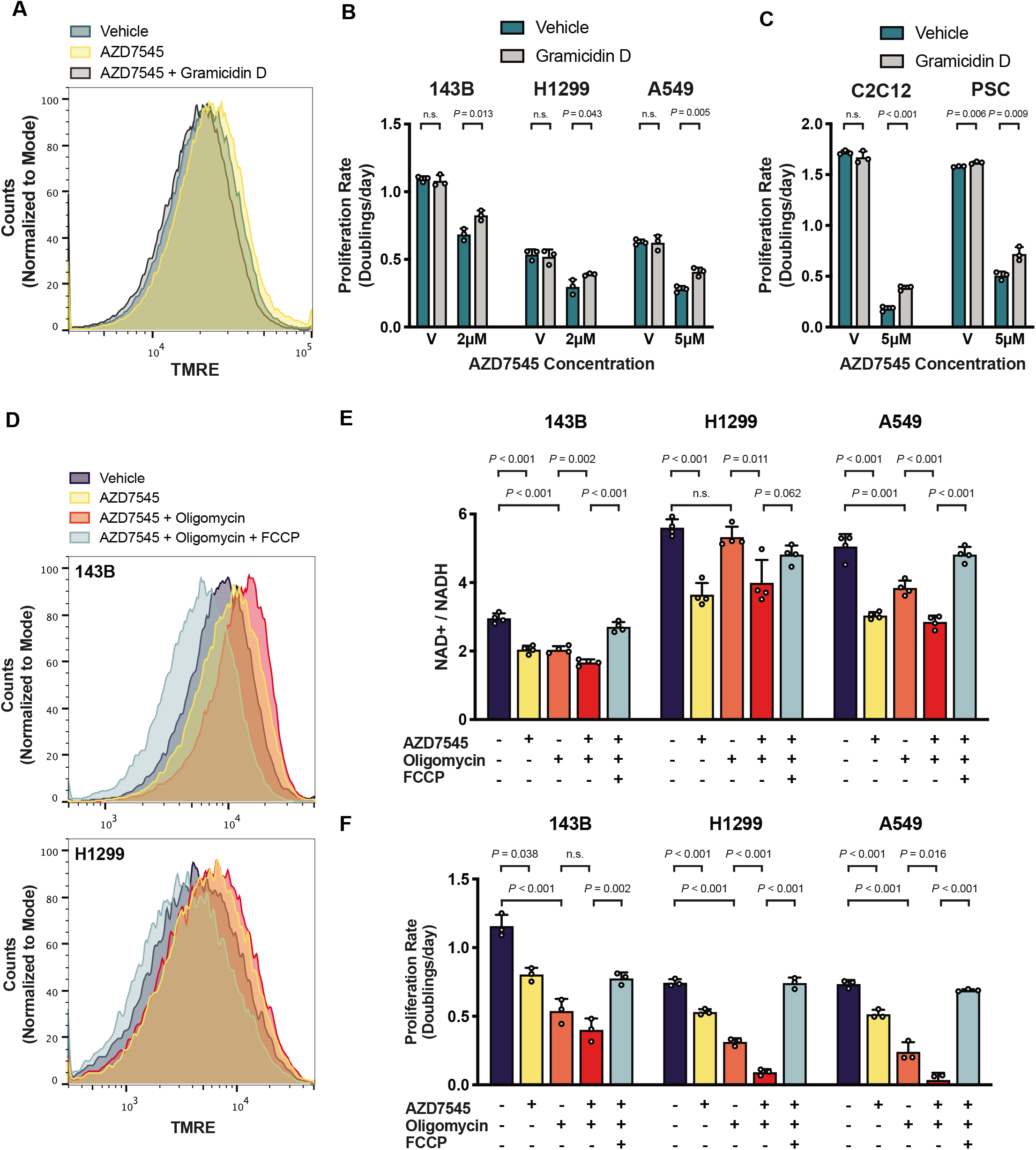
NAD+ regeneration by respiration is limited by the rate of mitochondrial ATP production. (A) Mitochondrial membrane potential, as reflected by TMRE fluorescence, of 143B cells cultured in vehicle, 2 μM AZD754, or 2 μM AZD754 with 5 nM gramicidin D for 5 hours. (B) Proliferation rate of 143B, H1299 and A549 cells treated with vehicle (V) or AZD7545 in the presence or absence of 0.5 nM gramicidin D (n = 3). (C) Proliferation rate C2C12 myoblasts or pancreatic stellate cells (PSC) that had been cultured with vehicle (V) or AZD7545 in the presence or absence of 1 nM gramicidin D (n = 3). (D) Mitochondrial membrane potential, as reflected by TMRE fluorescence, of 143B cells cultured for 5 hours with vehicle, 2 μM AZD7545, 1 nM oligomycin, and 1 μM FCCP as indicated. (E) NAD+/NADH ratio of 143B, H1299, and A549 cells cultured with vehicle, 5 μM AZD7545, 0.5 nM oligomycin, and 1 μM FCCP as indicated (n = 4). (F) Proliferation rate of 143B, H1299, and A549 cells cultured with vehicle, 5 μM AZD7545, 0.5 nM oligomycin and 1 μM FCCP as indicated (n = 3). Values shown in panel B, C, E, and F denote the mean ± SD. *P* values shown were calculated by unpaired, two-tailed Student’s t-test (n.s. = not significant).

### The demand for NAD+ regeneration can supersede the requirement for ATP in proliferating cells

To begin to test whether a mismatch in the demand for mitochondrial NAD+ regeneration relative to the demand for mitochondrial ATP production is what drives aerobic glycolysis, we utilized oligomycin, a selective inhibitor of the F_o_-subunit of the mitochondrial F_o_F_1_-ATP synthase. Oligomycin inhibits mitochondrial ATP production and reduces respiration by increasing ΔΨ (M. D. Brand & Nicholls, 2011). Culturing PDK-inhibited cells in the presence of oligomycin further increases ΔΨ beyond that observed with AZD7545 treatment alone (Figure 5D). Therefore, oligomycin is expected to further decrease the capacity for NAD+ regeneration via mitochondrial respiration. We assessed cell proliferation and NAD+/NADH ratio under these conditions, and found that oligomycin potentiates the proliferation defect and the NAD+/NADH reduction caused by PDK inhibition (Figure 5E,F). FCCP treatment, which collapses ΔΨ and prevents ATP synthesis by the mitochondrial F_o_F_1_-ATP synthase, restores the NAD+/NADH ratio and the proliferation of cells treated with both oligomycin and AZD7545. These data suggest that mitochondrial respiration to allow NAD+ regeneration can be more important than mitochondrial ATP production in some proliferating cells, including cells that otherwise engage in aerobic glycolysis.

The finding that uncoupling mitochondrial respiration from ATP production can rescue proliferation of PDK-inhibited cells argues that excess cell ATP can limit NAD+ regeneration. Consistent with the demand for NAD+ regeneration exceeding the demand for mitochondrial ATP synthesis to support proliferation in some cells that engage in aerobic glycolysis, we find FCCP can dose-dependently increase baseline cell proliferation in the absence of any other interventions (Figure 6A). These data suggest that excess ATP can impair NAD+ regeneration and proliferation even in standard culture conditions with abundant oxygen, and argue that coupling between mitochondrial respiration and ATP synthesis can constrain the extent to which respiration can support NAD+ regeneration and proliferation. Thus, cells may engage in aerobic glycolysis when the demand for NAD+ to support oxidation reactions is greater than the rate of ATP turnover.

**Figure 6.**
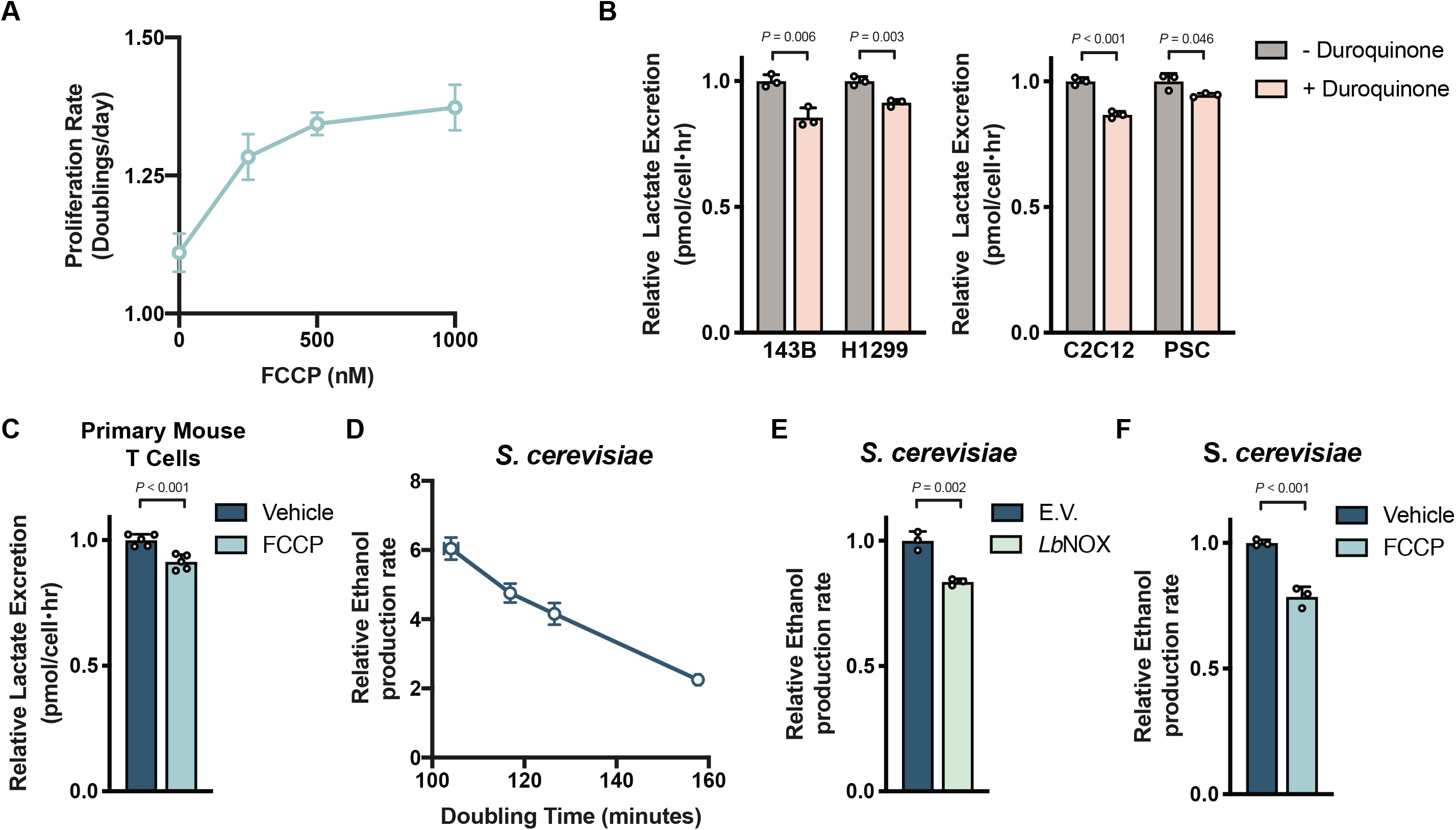
Aerobic glycolysis reflects cellular NAD+ availability. (A) The proliferation rate of 143B cells treated with the indicated concentration of FCCP (n = 3). (B) Relative lactate excretion of cells cultured in the presence or absence of duroquinone. The duroquinone concentration used was 4μM, 16μM, 8μM and 64μM for 143B, H1299, C2C12, and PSC cells, respectively (n = 3). (C) Relative lactate excretion of primary mouse T cells that had been stimulated with anti-CD3/CD28 antibodies in the presence or absence of 250 nM FCCP for 1 day (n = 5). (D) The relationship between ethanol production and proliferation rate for *S. cerevisiae* as determined by altering glucose concentration in the culture medium (See figure supplement 6C,D) (n = 3). (E) Relative ethanol production rate by *S. cerevisiae* expressing empty vector (E.V.) or *LbNOX* cultured in standard 3% glucose containing medium (n = 3). (F) Relative ethanol rate production by *S. cerevisiae* treated with vehicle or 2 μM FCCP cultured in standard 3% glucose containing medium (n = 3). Values shown in all panels denote the mean ± SD. *P* values shown were calculated by unpaired, two-tailed Student’s t-test (n.s. = not significant).

### Cellular NAD+ availability determines whether cells engage in aerobic glycolysis

If increased fermentation in proliferating cells is driven by a demand for NAD+ in excess of the demand for ATP, orthogonal pathways that promote NAD+ regeneration would be expected to suppress the degree to which cells engage in aerobic glycolysis despite the same (or higher) redox and ATP requirements to support proliferation. Indeed, duroquinone can suppress the rate of lactate excretion by both cancer cells and non-transformed cells (Figure 6B, figure supplement 6A) despite a similar proliferation rate (figure supplement 6B). *Lb*NOX expression did not affect lactate excretion in the cells studied, but increased pyruvate excretion as previously reported (Titov et al., 2016)(figure supplement 6C), which reflects decreased NAD+ regeneration by LDH. Additionally, FCCP administration decreased lactate excretion by activated mouse T cells (Figure 6C) without changing proliferation rate (figure supplement 6D). Thus, increasing NAD+ regeneration either by providing exogenous electron acceptors or by uncoupling mitochondrial respiration from ATP synthesis suppresses aerobic glycolysis without decreasing proliferation rate. These data support the hypothesis that demand for NAD+, rather than how cells produce ATP, is what drives aerobic glycolysis in these rapidly proliferating mammalian cells.

Aerobic glycolysis is linked to rapid proliferation across many biological contexts. To test whether a mismatch in the demand for NAD+ and ATP contributes to aerobic glycolysis in another system we considered *Saccharomyces cerevisiae,* a yeast where fermentation of glucose to ethanol accompanies rapid proliferation even in aerobic conditions (De Deken, 1966). We confirmed that glucose availability can affect proliferation rate and ethanol production in batch cultures of *S. cerevisiae,* even in aerated cultures (figure supplement 6E,F). We found a linear relationship between proliferation and fermentation rates (Figure 6D), confirming that aerobic glycolysis is correlated with proliferation in this yeast. To test whether fermentation is driven by the increased NAD+ demand of rapid proliferation in *S. cerevisiae,* we expressed *Lb*NOX and assessed the effect on both proliferation and aerobic glycolysis. *Lb*NOX expression decreased ethanol production without altering proliferation rate (Figure 6E, figure supplement 6G), arguing that demand for NAD+ also promotes aerobic glycolysis in this organism, consistent with previous reports (Vemuri, Eiteman, McEwen, Olsson, & Nielsen, 2007).

To test the hypothesis that elevated mitochondrial membrane potential limits the ability of respiration to support NAD+ regeneration in *S. cerevisiae,* we also assessed the effect of FCCP on both proliferation and aerobic glycolysis. In agreement with our hypothesis, we saw that FCCP administration decreased ethanol production rate without altering proliferation (Figure 6F, figure supplement 6H). The fact that FCCP reduces mitochondrial ATP production, and yet leads to reduction in fermentation as an alternative ATP production pathway without affecting proliferation, argues that ATP is also in excess in rapidly proliferating yeast. When coupled with the finding that the link between rapid proliferation and aerobic glycolysis is broken if cells are allowed alternative pathways of NAD+ regeneration, these observations suggest that demand for NAD+ in excess of demand for ATP drives aerobic glycolysis in diverse organisms across the kingdoms of life, regardless of whether lactate or ethanol are produced as the product of fermentation.

## Discussion

Aerobic glycolysis is observed across species ranging from prokaryotes to specific mammalian cell types, and yet a generalizable explanation for this phenotype has been lacking. The data from this study suggest that engagement in aerobic glycolysis reflects a metabolic state where the demand for NAD+ exceeds the demand for ATP to support cell function. Oxidation of pyruvate, rather than fermentation, increases the demand for mitochondrial respiration to regenerate NAD+. However, because mitochondrial NAD+ regeneration is coupled to mitochondrial ATP synthesis, when the demand for NAD+ exceeds the demand for ATP, insufficient supply of ADP to allow coupled respiration leads to increased ΔΨ and constrains further increases in mitochondrial respiration. Thus, ATP hydrolysis, which supplies ADP as substrate for mitochondrial ATP synthesis and dissipates ΔΨ produced by the mitochondrial ETC, imposes an upper limit on the rate of mitochondrial NAD+ regeneration regardless of oxygen availability. Therefore, if the requirement for NAD+ to fuel oxidation reactions is greater than the rate of ATP hydrolysis, pyruvate oxidation is limited and the more reduced NAD+/NADH ratio promotes fermentation even if oxygen is present.

Though the reactions that regenerate NAD+ do not directly provide biomass to cells, this cofactor is needed to catabolize reduced nutrients, including sugars and lipids, and to synthesize oxidized macromolecules, such as nucleotides and amino acids (Hosios & Vander Heiden, 2018). Previous work has identified aspartate synthesis as a major NAD+ demand (Birsoy et al., 2015; Garcia-Bermudez et al., 2018; Gui et al., 2016; Sullivan et al., 2015; Sullivan et al., 2018), and cells require NAD+ to support additional cellular processes, including serine and folate synthesis (Bao et al., 2016; Diehl, Lewis, Fiske, & Vander Heiden, 2019), histone deacetylation by sirtuins, maintenance of calcium homeostasis, and poly(ADP-ribose) polymerase activity (Canto, Menzies, & Auwerx, 2015; Ying, 2008). Electron acceptor availability can restrict cell proliferation, and the NAD+/NADH ratio has been found to correlate with tumor growth in vivo (Gui et al., 2016). Non-proliferating cells with high anabolic demands, such as mammalian retina pigmented epithelial cells also require NAD+, and retinal toxicity has proven to be a liability for drugs that target NAD+ synthesis (Zabka et al., 2015).

Several explanations have been proposed for the Warburg effect, many of which consider how inefficient ATP production by aerobic glycolysis is sufficient to support the high ATP demands of proliferation. The finding that uncoupling respiration from ATP production allows increased pyruvate oxidation and continued proliferation, even in the presence of oligomycin, argues strongly that the requirement for NAD+ can be greater than the requirement for ATP in at least some proliferating cells. Furthermore, the observation that increasing ATP consumption also allows proliferation, despite increased pyruvate oxidation, suggests insufficient ATP consumption can limit mitochondria respiration in cells and that aerobic glycolysis may therefore reflect a state of high NAD+ demand and excess ATP. The idea that ATP hydrolysis, rather than ATP synthesis, could be limiting for cancer cell metabolism has historical support (Racker, 1972; Scholnick, Lang, & Racker, 1973), although this possibility is not considered by most studies. Nevertheless, it has been reported that some proliferating cells engage in futile metabolic cycles that consume ATP, such as fatty acid synthesis and oxidation (Yao et al., 2016), increased protein turnover (Y. Zhang et al., 2014), or the creatine shuttle (Kurmi et al., 2018) for unclear reasons. These futile cycles may promote ATP consumption to support oxidized biomass synthesis, and could explain why increased ATP turnover involving ENTPD5 can promote the proliferation of Pten-null cancers or why increasing the ATP/AMP ratio can result in tumor regression (Fang et al., 2010; Naguib et al., 2018). The mechanism by which ATP consumption promotes proliferation in those contexts remains unexplained, but a model where excess ATP limits NAD+ regeneration by respiration is consistent with increasing ATP consumption enabling proliferation.

While a demand for NAD+ that exceeds the demand for ATP can explain why some cells engage in fermentation rather than oxidative metabolism, this model does not explain why increased glucose uptake and glycolysis are also associated with proliferation. Fermentation is redox neutral and does not net regenerate NAD+. In fact, the only known output of fermentation is ATP, and if excess ATP limits net NAD+ regeneration to promote fermentation, then why glucose metabolism is increased remains unexplained. Nevertheless, a model where aerobic glycolysis reflects a metabolic state in which the requirement for NAD+ supersedes the demand for ATP is consistent with the reality of the Warburg effect, which is best characterized by a relative increase in fermentation, rather than a complete switch from mitochondrial respiration to glycolysis. It also fits with the broad association between aerobic glycolysis and proliferation (Diaz-Ruiz, Rigoulet, & Devin, 2011), the link between this phenotype and nucleotide synthesis (Lunt et al., 2015; Wang et al., 1976), the continued dependence of most proliferating cells on respiration (Howell & Sager, 1979; Tan et al., 2015; Weinberg et al., 2010; Wheaton et al., 2014; X. Zhang et al., 2014), and that fact that reversing aerobic glycolysis slows rather than stops proliferation.

Different species of yeast diverge in their response to glucose in terms of their propensity to ferment carbon (De Deken, 1966), and organisms that proliferate rapidly without engaging in aerobic glycolysis may have evolved mechanisms to prevent a mismatch in the demand for NAD+ and ATP. Of note, regulated uncoupling of mitochondrial respiration from ATP synthesis is important for thermoregulation and these same processes could also facilitate rapid proliferation. While mammalian uncoupling proteins have limited tissue expression and are tightly regulated, these proteins as well as other means to uncouple mitochondrial respiration (Bertholet et al., 2019) may support NAD+ regeneration when NAD+ demand exceeds the demand for ATP in some contexts. For example, lymphocytes can proliferate faster than most other mammalian cells, and may use such an adaptive program to support proliferation, potentially explaining why we observe variability in whether FCCP-mediated mitochondrial uncoupling can enhance T cell proliferation. However, the fact that maintaining a high ATP/ADP ratio is essential for survival of all cells may explain why nutrient oxidation remains tightly coupled in ATP production in most cells and leads to aerobic glycolysis being engaged when the demand for NAD+ exceeds the demand for ATP in cells.

## Materials and Methods

### Cell Culture Experiments

Cell lines were maintained in DMEM (Corning 10-013-CV) supplemented with 10% fetal bovine serum. For all experiments, cells were washed three times in phosphate buffered saline (PBS), and then cultured in DMEM without pyruvate (Corning 10-017-CV) with 10% dialyzed fetal bovine serum, supplemented with the indicated treatment condition. The reagents used for the cell culture experiments are as follows: AZD7545 (Selleck Chemicals, s7517), sodium pyruvate (Sigma, P2256), sodium L-lactate (Sigma, L7022), duroquinone (Sigma, D223204), metformin hydrochloride (Sigma PHR1084), FCCP (carbonyl cyanide 4- (trifluoromethoxy)phenylhydrazone, also referred to as trifluoromethoxy carbonylcyanide phenylhydrazone) (Sigma, C2920), oligomycin A (Sigma 75351), gramicidin from *Bacillus aneurinolyticus (Bacillus brevis)* (Sigma, G5002). All cells were cultured at 37°C with 5% CO_2_.

### CRISPRi/Cas9-mediated repression of PDK1

PDK1 expression was suppressed using CRISPR interference (CRISPRi)-mediated transcriptional repression. A CRISPRi library (Horlbeck et al., 2016) was used to design sgRNA human targeting PDK1 (Guide 1: F 5’GCTCACGTACCACTCGGCAG 3’; Guide 2: F 5’GACGTCCCTCACGTACCACT 3’) or a non-targeting control (NTC; F 5’GGGAACCACATGGAATTCGA 3’). The sgRNA were cloned into a modified LentiCRISPRv2 plasmid, in which Cas9 was mutated to nuclease-dead Cas9 and fused to repressive chromatin modifier domain KRAB (Krüppel-associated box) domain of Kox1 (dCAS9-KRAB) (Gilbert et al., 2013). A pooled population of stable knockdown cell lines were generated and maintained in 5μM/mL Blasticidin.

### Pancreatic Stellate Cell (PSC) isolation

PSCs were isolated from β-actin-GFP mice (The Jackson Laboratory, 006567) in which 3 mL of 1.3 mg/mL cold collagenase P (Sigma 11213865001) and 0.01 mg/mL DNAse (Sigma D5025) in GBSS (Sigma G9779) were injected into the pancreas. The tissue was then placed into 2 mL of 1.3 mg/mL collagenase P solution on ice. Cells were then placed in a 37°C water bath for 15 minutes. The digested pancreas was filtered through a 250 μm strainer and washed with GBSS with 0.3% BSA. A gradient was created by suspending the cells in Nycodenz (VWR 100356-726) and layering in GBSS with 0.3% BSA. Cells were then centrifuged at 1300×g for 20 minutes at 4°C. The layer containing PSCs was removed, filtered through a 70 μm strainer, washed in GBSS with 0.3% BSA, and plated for cell culture in DMEM with 10% FBS and penicillin-streptomycin. Post-isolation, PSCs were immortalized with TERT and LargeT Antigen overexpression.

### T cell isolation and culture

Primary human CD4+ T cells were isolated from peripheral blood by density gradient centrifugation over Lympholyte-H (Cedarlane Laboratories) and negative selection using the Dynabeads Untouched Human CD4 T Cells kit (Invitrogen 11346D) according to the manufacturer’s instructions. Purity was assessed by flow cytometry for CD3 and CD4 and routinely found to be ≥95%. Cells were activated using Dynabeads Human T-Activator CD3/CD28 beads (Invitrogen) according to the manufacturer’s instructions and cultured in RPMI supplemented with 10% FBS and 30 U/ml recombinant human IL-2 (PeproTech) at 37°C in 5% CO_2_. Primary mouse T cells were isolated from the spleens and lymph nodes of C57BL/6J mice using a Pan-T cell isolation kit (MACS 130-095-130). Isolated murine T cells were activated on plates coated with anti-CD3e (BD Biosciences 553057) and anti-CD28 (BD Biosciences 553294) antibodies and cultured in RPMI with 10% FBS (non-dialyzed), 1% P/S, 1x NEAA culture supplement (Thermo Fischer Scientific, Cat. #: 11140050), and 50μM β-mercaptoethanol.

### T cell proliferation assay

Analysis of primary human CD4+ T cell and primary mouse T cell proliferation by dye dilution was carried out as described previously (Matheson et al., 2015). Freshly isolated cells were incubated at 1 × 10^7^ cells/ ml in PBS with 0.5% dialyzed FBS supplemented with 5μM carboxyfluorescein succinimidyl ester (CFSE, Molecular Probes) or 2.5μM CellTrace Far Red (Molecular Probes) for 5-7 minutes at room temperature. Staining was quenched with ice-cold complete media and washed cells were resuspended in RPMI supplemented with 10% dialyzed FBS and the indicated treatment condition before activation. For human CD4+ T cells, cells were expanded in fresh media containing the indicated treatment condition and CFSE fluorescence was then measured by flow cytometry after another 2 days. Primary murine T cells were expanded in fresh media containing the indicated treatment condition for 2 days, and CFSE fluorescence was then measured by flow cytometry.

### Flow Cytometry of T cells

CD3/CD28 Dynabeads were removed from stimulated primary human CD4+ T cells using a DynaMag-2 magnet (Invitrogen). For antibody staining, cells were incubated for 30 minutes at 4°C in PBS with a fluorochrome-conjugated antibody (mouse monoclonal APC-conjugated anti-CD69 (BioLegend 10909); mouse monoclonal PE-conjugated anti-CD25 (Pharmingen 30795X) as indicated). For antibody staining or fluorescent dye dilution, cells were analyzed using a FACSCanto II, FACSCalibur or LSR Fortessa (BD Biosciences) flow cytometer. Data processing (including proliferation analysis) was conducted using FlowJo V10.6.1. For mouse T cell antibody staining, cells were incubated for 30 minutes at 4°C in PBS containing 2% fetal bovine serum with eFluor780 viability dye (eBioscience 65-0865-18) and fluorochrome-conjugated antibodies: mouse monoclonal PE-CF594-conjugated anti-CD45 (BD Horizon 562420) and eFluor450-conjugated anti-CD3e (Thermo Fisher Scientific 48-0032-80); TMRE staining (Thermo Fisher Scientific T6690) was performed at room temperature in FluoroBrite DMEM (Thermo Fischer Scientific A1896701) supplemented with dialyzed FBS. Stained cells were analyzed using a LSR-II or LSR Fortessa (BD Biosciences) flow cytometer. Data processing (including proliferation analysis) was conducted using FlowJo V10.6.1.

### Generation of LbNOX-Expressing Cells

LbNOX-FLAG cDNA was cloned from pUC57-LbNOX using the primers 5’-GGGGACAAGTTTGTACAAAAAAGCAGGCATGAAGGTCACCGTGGTC-3’ and 5’-GGGGACCACTTTGTACAAGAAAGCTGGGTTTACTTGTCATCGTCATCCTTGTAATC-3’. pUC57-LbNOX was a gift from Vamsi Mootha (Addgene, 75285). The PCR product was subsequently cloned into pInducer20 using LR Clonase II Plus (ThermoFisher 12538120). pInducer20 was a gift from Stephen Elledge (Addgene, 44012). A549, 143B, and H1299 cells were transduced with lentivirus containing pInducer20-LbNOX-FLAG or pInducer20-E.V. and 10ug/mL polybrene (Millipore TR-1003-G). The infected cells were selected in 1 mg/mL G418 (VWR-G5005). For proliferation experiments done with these cell lines, all conditions were supplemented with 500 ng/mL doxycycline.

### Proliferation Rates

Proliferation rate was determined as previously described (Gui et al., 2016; Sullivan et al., 2015). Briefly, cells were plated in replicate six-well plates in 2 mL at an initial seeding density of 20,000 cells per well for all cells except for MDA-MB-231, which were seeded at 40,000 cells per well. Cells were permitted to settle overnight and one six-well dish was counted to calculate the starting cell number at the initiation of the experiment. For all remaining dishes, cells were washed three times with PBS and 4 mL of treatment media was added to each well. After 48 hours, cells were washed again three times with PBS and 4 mL of the treatment media was replenished. Four days after the initial treatment, cells were counted to obtain the final cell counts for the experiment. Counts were done using a Cellometer Auto T4 Plus Cell Counter (Nexcelcom Bioscience) or by sulforhodamine (SRB) assay. For cell quantification by SRB, cells were fixed by adding 10% trichloroacetic acid and incubated 4°C for at least one hour. Fixed cells were washed with deionized water and then stained with 0.057% sulforhodamine B in 1% acetic acid for 30 min. Following three washes with 1% acetic acid, plates were air-dried at room temperature. To solubilize the SRB dye, 1 mL of 10 mM Tris [pH 10.5] was added per well, and absorbance was measured at 510 nm using a microplate reader (Tecan Infinite M200Pro). The proliferation rate was calculated as follows.

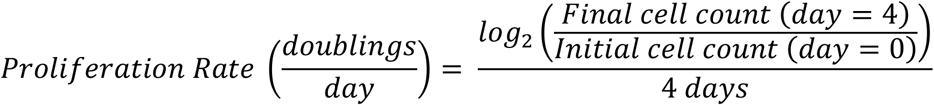

### LCMS Metabolite Measurement

Metabolites were measured by LCMS on a QExactive bench top orbitrap mass spectrometer equipped with an Ion Max source and a HESI II probe, which was coupled to a Dionex UltiMate 3000 HPLC system (Thermo Fisher Scientific, San Jose, CA). External mass calibration was performed using the standard calibration mixture every 7 days. For each sample, 4 μL of each sample was injected onto a SeQuant^®^ ZIC^®^-pHILIC 150 × 2.1 mm analytical column equipped with a 2.1 × 20 mm guard column (both 5 mm particle size; EMD Millipore). Buffer A was 20 mM ammonium carbonate, 0.1% ammonium hydroxide; Buffer B was acetonitrile. The column oven and autosampler tray were held at 25°C and 4°C, respectively. The chromatographic gradient was run at a flow rate of 0.150 mL/min as follows: 0-20 min: linear gradient from 80-20% B; 20-20.5 min: linear gradient form 20-80% B; 20.5-28 min: hold at 80% B. The mass spectrometer was operated in full-scan, polarity-switching mode, with the spray voltage set to 3.0 kV, the heated capillary held at 275°C, and the HESI probe held at 350°C. The sheath gas flow was set to 40 units, the auxiliary gas flow was set to 15 units, and the sweep gas flow was set to 1 unit. MS data acquisition was performed in a range of m/z = 70–1000, with the resolution set at 70,000, the AGC target at 1 × 10^6^, and the maximum injection time (Max IT) at 20 msec. For detection of ^13^C-labeled citrate, targeted selected ion monitoring (tSIM) scans in negative mode centered on 194.1985 was included. The isolation window was set at 8.0 m/z. For all tSIM scans, the resolution was set at 70,000, the AGC target was 1105, and the max IT was 250 ms. Relative quantitation of polar metabolites was performed with XCalibur QuanBrowser 2.2 (Thermo Fisher Scientific) using a 5 ppm mass tolerance and referencing an in-house library of chemical standards.

### GCMS Metabolite Measurement

Gas-chromatography coupled to mass spectrometry (GCMS) analysis was done as described previously (Lewis et al., 2014). Dried metabolite samples were derivatized with 20 μL of methoxamine (MOX) reagent (ThermoFisher TS-45950) and 25 μL of N-tert-butyldimethylsilyl-N-methyltrifluoroacetamide with 1% tert-butyldimethylchlorosilane (Sigma 375934). Following derivatization, samples were analyzed using a DB-35MS column (30m × 0.25mm i.d. × 0.25 μm, Agilent J&W Scientific) in an Agilent 7890 gas chromatograph (GC) coupled to an Agilent 5975C mass spectrometer (MS). Data were corrected for natural isotope abundance using inhouse algorithms as in (Lewis et al., 2014).

### Dynamic stable-isotope labeling experiments

Cells were plated in six-well plates at a seeding density of 150,000 cells per well and cultured overnight. Prior to the initiation of the experiment, cells were washed three times with PBS, and then cultured in 1.5 mL media containing 5mM glucose and the indicated treatment condition prior to tracing glucose fate. To assess glucose fate, 30 μL of a 1 M [U-^13^C_6_]glucose solution was added to a final concentration of 20 mM for the indicated time (ranging from 2 to 12 minutes). At each time point wells were washed as quickly as possible with ice-cold blood bank saline and lysed on the dish with 300 μL of ice-cold 80% HPLC grade methanol in HPLC grade water. Samples were scraped, collected into Eppendorf tubes, and vortexed for 10 minutes at 4°C. Samples were centrifuged at 21,000×g for 10 minutes at 4°C to precipitate protein. 50 μL of each sample was collected for immediate analysis by LCMS, or the supernatant was dried down under nitrogen gas for subsequent analysis by GCMS.

### Oxygen Consumption

Oxygen consumption rates (OCR) was measured using an Agilent Seahorse Bioscience Extracellular Flux Analyzer (XF24) using standard methods. Briefly, cells were plated at 50,000 cells per well in Seahorse Bioscience 24-well plates in 50 μL of DMEM without pyruvate (Corning 10-017-CV) supplemented with 10% dialyzed fetal bovine serum. An additional 500μL of media was added following a one hour incubation. The following day, cells were washed three times with PBS and incubated in DMEM without pyruvate with the indicated treatment. Five hours later, OCR measurements were made every 6 minutes, and injections of pyruvate at 16 minutes, FCCP (Sigma, C2920) at 32 minutes, and rotenone (Sigma R8875, 2 μM) and antimycin (Sigma A8674, 2 μM) at 48 minutes. Basal OCR was calculated by subtracting residual OCR following the addition of rotenone and antimycin from the initial OCR measurements.

### Glucose/Lactate Excretion Index

Medium was collected from cells cultured in vehicle or 0.25 μM AZD7545 for 48 hours. Glucose and lactate concentrations were measured on a YSI-2900 Biochemistry Analyzer as described previously, and rate of lactate excretion into media was normalized to consumption of glucose (Hosios et al., 2016).

### Lactate and Pyruvate Excretion

Medium was collected from 143B cells cultured in pyruvate-free DMEM treated with vehicle or 200μM duroquinone for 48 hours, or medium was collected from 143B cells cultured in pyruvate-free DMEM expressing empty vector or *Lb*NOX for 48 hours. Extracellular pyruvate and lactate were measuring using GCMS, using a labeled internal standard (Cambridge Isotopes CLM-1579-PK), and the excretion was normalized to the proliferation rate to calculate the rate per unit growth. For H1299, PSC, and C2C12 cells, extracellular lactate was measured using a YSI-2900 Biochemistry Analyzer after treatment with the indicated amount of duroquinone for 48 hours in pyruvate-free DMEM. Lactate excretion in primary mouse T cells was measured using a YSI-2900 Biochemistry Analyzer after 24 hours of stimulation using anti-CD3e (BD Biosciences 553057) and anti-CD28 (BD Biosciences 553294) antibodies and cultured in RPMI with 10% FBS, 1% P/S, 1x NEAA culture supplement (Thermo Fischer Scientific, Cat. #: 11140050), and 50μM β-mercaptoethanol. Lactate excretion was normalized to proliferation rate as assessed by counting the cells using a Cellometer Auto T4 Plus Cell Counter (Nexcelcom Bioscience).

### Western Blot Analysis

Cells washed with ice-cold PBS, and scraped into cold RIPA buffer containing cOmplete Mini protease inhibitor (Roche 11836170001) and PhosStop Phosphatase Inhibitor Cocktail Tablets (Roche 04906845001). Protein concentration was calculated using the BCA Protein Assay (Pierce 23225) with BSA as a standard. Lysates were resolved by SDS-PAGE and proteins were transferred onto nitrocellulose membranes using the iBlot2 Dry Blotting System (Thermo Fisher, IB21001, IB23001). Protein was detected with the primary antibodies anti-Pyruvate Dehydrogenase E1-alpha subunit (phospho S293) (Abcam, ab92696), anti-PDK1 (Cell Signaling Technologies, 3062S), anti-FLAG (Sigma, F1804) and anti-Vinculin (Sigma, V9131). The secondary antibodies used were IR680LT dye conjugated anti-rabbit IgG (Licor Biosciences 925-68021), IRDye 800CW conjugated anti-mouse IgG (Licor Biosciences 925-32210), HRP-conjugated anti-rabbit IgG (Millipore 12–348), and HRP-conjugated anti-mouse IgG (Millipore 12-349).

### Tumor Growth in Mice

Two million A549 or one million 143B cells were injected into the flanks of 4-6 week old, male NU/NU mice (Charles River Laboratories, 088). A caliper was used to measure flank tumor volume in two dimensions and volume was calculated using the equation V = (π/6)(L×W^2^). Length was defined to be the longer of the two dimensions measured. For the A549 xenografts, the tumors were permitted to reach a size of 50 mm^3^, after which the animals were randomly assigned to an experimental group and the treatment regimen was initiated. Metformin (500 mg/kg) and AZD7545 (45 mg/kg) were both administered once daily by oral gavage, with a vehicle of water and 0.5% (w/w) methocel/0.1% polysorbate 80 respectively. All animal experiments were approved by the MIT Committee on Animal Care.

### NAD+/NADH Measurements

Cells were seeded at 20,000 cells per well in six-well plates and permitted to adhere overnight. Cells were then washed three times in PBS and incubated in 4 mL of the indicated treatment media for 5 hours prior to rapidly washing three times in 4°C PBS and extraction in 100μL of ice-cold lysis buffer (1% dodecyltrimethylammonium bromide [DTAB] in 0.2 N of NaOH diluted 1:1 with PBS), then snap-frozen in liquid nitrogen and frozen at −80°C. The NAD+/NADH ratio was measured using an NAD/NADH-Glo Assay kit (Promega G9072) according to a modified protocol as described previously (Gui et al., 2016). Briefly, to measure NAD+, 20 μL of lysate was transferred to PCR and diluted with 20 μL of lysis buffer and 20 μL 0.4 N HCl, and subsequently incubated at 60°C for 15 minutes. For NADH measurement, 20 μL of freshly thawed lysate was transferred to PCR tubes and incubated at 75°C for 30 minutes. The acidic conditions permit for selective degradation of NADH, while the basic conditions degrade NAD+. Following the incubation, samples were spun on a bench-top centrifuge and quenched with 20 μL neutralizing solution. The neutralizing solution consisted of 0.5 M Tris base for NAD+ samples and 0.25 M Tris in 0.2 N HCl for the NADH samples. The instructions in the Promega G9072 technical manual were then followed to measure NAD+ and NADH levels using a luminometer (Tecan Infinite M200Pro).

### Yeast proliferation and metabolite analysis

In studies comparing the effects of *Lb*NOX on ethanol production, CEN.PK 5D (MATa ura3-52 HIS3, LEU2 TRP1 MAL2-8c SUC2) expressing p416-TEF-lbNOX or p416-TEF (empty vector) were grown in log phase in YPD (3% glucose) at 30°C. At OD = 0.8, supernatant was collected and analyzed for ethanol content using an Ethanol Assay Kit (Sigma, MAK706). Rate of ethanol production was normalized to the proliferation rate. To study the effects of FCCP on ethanol production, wild type CEN.PK 5D strain yeast were cultured in YPD (3% glucose) at 30°C in the indicated amount of FCCP.

### Mitochondrial Membrane Potential Measurement

Mitochondrial membrane potential was assessed using the TMRE (tetramethylrhodamine, ethyl ester) assay kit (Abcam, ab113852) in nonquench mode. The concentration of TMRE in nonquench mode was determined empirically (Perry et al., 2011). Cells were plated in 6 well dishes at a plating density of 150,000 cells per well and incubated in media containing the indicated concentration of vehicle AZD7545, 500 nM FCCP, 1 nM oligomycin, and/or 5nM gramicidin D. The cells were then treated with 200 nM TMRE for 30 minutes, trypsinized, washed with PBS, and resuspended in Gibco FluoroBrite DMEM (Thermo Fisher Scientific A1896701) supplemented with 10% dialyzed FBS. TMRE fluorescence was measured on a BD LSR II flow cytometer.

## Acknowledgements

We thank all members of the Vander Heiden lab for thoughtful discussion and advice. We also thank Allison N. Lau for isolation of pancreatic stellate cells. This work was supported by the Ludwig Center for Molecular Oncology Fund (A.L.), NSF GRFP DGE-1122374 (A.L.), NIH T32GM007287 (A.L.; Z.L.; D.Y.G.; M.Z.), NIH T32GM007753 (B.T.D.), NIH K99CA218679 (L.B.S.), HHMI Medical Research Fellows Program (A.A.), MRC CSF MR/P008801/1 (N.J.M.), NHSBT WPA15-02 (N.J.M), the Novo Nordisk Foundation Grant No. NNF10CC1016517 (R.F.) and the Knut and Alice Wallenberg Foundation (R.F.). M.G.V.H. acknowledges support from a Faculty Scholar grant from the Howard Hughes Medical Institute, SU2C, the Lustgarten Foundation, the MIT Center for Precision Cancer Medicine, the Ludwig Center at MIT, and the NIH (R35CA242379, R01CA201276, R01CA168653, P30CA14051). We also thank The Swanson Biotechnology Center Flow Cytometry Facility.

## Author Contributions

A.L., Z.L., D.Y.G., L.B.S., M.Z., A.N, A.A., and N.J.M. performed the experiments. B.T.D. provided computational support. C.J.T. provided critical supplies and experimental design. C.A.L. performed the targeted LCMS experiments. R.F. generated the *Lb*NOX expressing yeast strain. S.S. and N.J.M. provided immunology advice. A.L., Z.L. and M.G.V.H. designed the study and wrote the manuscript with input from all authors.

## Competing Interests

The authors are aware of no competing interests; however, M.G.V.H. discloses he is a scientific advisor for Agios Pharmaceuticals, Aeglea Biotherapeutics, Auron Therapeutics, and iTeos Therapeutics. S.S. is member of the scientific advisory board of Arcus Biosciences, Venn Therapeutics, Tango Therapeutics, Replimune and serves as a scientific advisor for Dragonfly Therapeutics, Merck, Ribon, Torque and TAKEDA. A.L. is a current employee of a Flagship Pioneering biotechnology start-up company.

