## Supplemental Text for "Increased demand for NAD+ relative to ATP drives aerobic glycolysis"

**Figure supplement legends:**

**Figure supplement 1. PDK inhibition increases oxygen consumption and slows both cell proliferation and tumor growth.**

(A) Kinetic labeling of citrate from $^{13}$C-labeled glucose to assess PDH flux with and without PDK inhibition by AZD7545. 143B cells were incubated for 1 hour in media containing 5 mM unlabeled glucose with vehicle or 5 μM AZD7545, after which 20 mM [U-$^{13}C_6$]glucose was added. The fraction of M+2 citrate was measured by LCMS following the addition of $^{13}$C-labeled glucose (n = 3 per time point).

(B) Oxygen consumption rate (OCR) measured in HeLa, 143B, and H1299 cells treated with vehicle or 5 μM AZD7545 for 5 hours, with the addition of 1 mM pyruvate, 1.5 μM of the mitochondrial uncoupler trifluoromethoxy carbonylcyanide phenylhydrazone (FCCP), and 1.5 μM each of the complex I inhibitor rotenone and the complex III inhibitor antimycin A as indicated. Basal OCR in the absence of pyruvate or mitochondrial inhibitors is presented in Figure 1D (n = 12).

(C) The percent of primary human CD4+ T cells that had divided at least once over a period of 4 days in the presence of vehicle or 5 μM AZD7545. Cell division was assessed by CFSE or CellTrace Far Red dilution measured by flow cytometry 4 days following cell stimulation with CD3/CD28 dynabeads in the indicated treatment condition. Three biological replicates of primary human CD4+ T cells collected from different donors and analyzed as independent experiments were assessed. Representative data are presented in Figure 1H. Conditions were compared using a paired 2-tailed Student's t-test (n = 3).

(D) The percent of primary murine T cells that had divided at least once over a period of 2 days in the presence of vehicle or 5 μM AZD7545. CFSE fluorescence was assessed by flow cytometry 2 days following stimulation with anti-CD3/CD28 antibodies in the presence of the indicated treatment condition. Representative data are presented in Figure 1I (n = 8). Conditions were compared using an unpaired 2-tailed Student's t-test

(E) Primary human CD4+ T cells were isolated and stimulated as described in (C) in the presence of vehicle or 5 μM AZD7545 prior to staining with fluorochrome-conjugated anti-CD69 or anti-CD25 antibodies and analyzed by flow cytometry. The fold change (log2) in geometric mean fluorescence intensity compared with resting cells is shown for each of these markers of T cell activation from three biological replicates, as is representative data from one donor. Unstained, stimulated cells (dotted black line) and stained, unstimulated cells (resting, grey) are also shown as controls.

(F) Western blot analysis to assess expression of pyruvate dehydrogenase kinase 1 (PDK1) in 143B cells that were transduced with sgRNA targeting PDK1 or a non-targeting control (NTC). Expression of vinculin was also assessed as a loading control.

(G) Growth of tumors arising in nude mice following injection of the cells described in (F) (n = 15).

For panels A, B, C, D, E and G, values denote mean ± SEM.

**Figure supplement 2. PDK inhibition decreases the NAD+/NADH ratio and slows cell proliferation.**

(A) The NAD+/NADH ratio of A549 and HeLa cells treated with vehicle or 5 μM AZD7545 for 5 hours (n = 4).

(B) Proliferation rate of A549 and HeLa cells treated with vehicle (V) or AZD7545 in the presence or absence of 1 mM pyruvate (n = 3).

(C) Proliferation of primary human CD4+ T cells in vehicle or increasing concentrations of AZD7545 in the presence (blue) or absence (orange) of 1 mM pyruvate. Human CD4+ T cells were isolated and stained with CFSE prior to stimulation with CD3/CD28 Dynabeads and CFSE fluorescence was assessed by flow cytometry after 4 days. Data from CFSE-stained, unstimulated cells (grey) that did not proliferate are also shown as controls.

(D) The percent of the original human CD4+ T cell population that has divided at least once in the presence of vehicle or AZD7545 with or without 1 mM pyruvate. Cell division was assessed by CFSE or CellTrace Far Red dilution measured by flow cytometry 4 days following cell stimulation with CD3/CD28 dynabeads in the indicated treatment condition. Representative data are shown in panel (C) and in Figure 2G. *P* values were calculated by paired, two-tailed Student's t-test on three biological replicates from different donors (n = 3).

(E) Proliferation of primary mouse T cells in the presence of vehicle or AZD7545 with (blue) or without (orange) 1 mM pyruvate as indicated. CFSE fluorescence was assessed by flow cytometry 2 days following stimulation with anti-CD3/CD28 antibodies in the presence of the indicated treatment condition. Data from CFSE-stained unstimulated cells (grey) are also shown as controls.

(F) The percent of the original mouse T cell population that divided at least once in the presence of vehicle or AZD7545 with or without 1 mM pyruvate. CFSE fluorescence was assessed by flow cytometry 2 days following stimulation with anti-CD3/CD28 antibodies in the presence of the indicated treatment condition. Representative data are presented in panel (E) and in Figure 2H.

(G) The NAD+/NADH ratio of HeLa cells cultured in media containing vehicle, 5 μM AZD7545, or 5 μM AZD7545 and 10 mM lactate (n = 4).

(H) Proliferation rate of HeLa cells treated with vehicle (V) or AZD7545 and supplemented with 1 mM or 10 mM lactate as indicated (n = 3).

(J) Proliferation rate of 143B and H1299 cells cultured in vehicle or 5 μM AZD7545 without pyruvate or lactate (-), or in media containing the indicated ratio of lactate to pyruvate where lactate was present at 10mM in all cases except for at the ratio labeled "0". For the condition where the ratio is labeled "0", lactate was not added and the pyruvate concentration was 1 mM. For panels A, B, D, F, G, H, and I values shown denote the mean ± SD. *P* values were calculated by unpaired, two-tailed Student's t-test unless otherwise indicated (n.s. = non-significant).

**Figure supplement 3. Interventions that increase the NAD+/NADH ratio reduce sensitivity to PDK inhibition.**

(A) Proliferation rate of A549 cells cultured with vehicle (V) or AZD7545 with or without 20 μM duroquinone as indicated (n = 3).

(B) Western blot analysis using an anti-FLAG antibody to examine expression of *Lb*NOX in 143B, A549, and HeLa cells that had been infected with empty vector (E.V.) or doxycycline inducible FLAG-tagged *Lb*NOX (FLAG-*Lb*NOX) and cultured in the presence of 500 ng/mL doxycycline. Expression of vinculin was also assessed as a loading control.

(C) Proliferation rate of A549 cells expressing empty vector (E.V.) or *Lb*NOX in the presence of vehicle (V) or AZD7545 as indicated. All cells were cultured with doxycycline (500 ng/mL) to induce *Lb*NOX expression (n = 3).

(D) Relative aspartate levels, normalized to the vehicle-treated condition, as measured by GCMS in cells cultured in presence of vehicle or 5 μM AZD7545 for 5 hours (n = 4).

(E) Relative aspartate levels, normalized to the no pyruvate condition, as measured by GCMS in cells treated with 5 μM AZD7545 in the absence or presence of 1 mM pyruvate (n = 3).

(F) Relative aspartate levels, normalized to the vehicle-treated condition, as measured by LCMS in 143B cells treated with 5 μM AZD7545 with or without 20 μM duroquinone (n = 3).

(G) Relative aspartate levels, normalized to the E.V. vehicle-treated condition, as measured by LCMS in 143B cells expressing empty vector (E.V.) or *Lb*NOX cultured in the presence of 5 μM AZD7545 and 500 ng/mL doxycycline (n = 3).

(H) Relative citrate levels, normalized to the vehicle-treated condition, as measured by LCMS in 143B cells treated with vehicle or 5 μM AZD7545 (n = 6).

(I) Relative palmitate levels, normalized to the vehicle-treated condition, as measured by LCMS in 143B cells treated with vehicle or in 5 μM AZD7545 (n = 6).

(J) Intracellular metabolites, as measured by LCMS, of 143B cells treated with vehicle or 5 μM AZD7545 in the presence or absence of 20 μM duroquinone. Metabolite levels were compared to the vehicle-treated condition, and the fold change of all metabolites were hierarchically clustered and displayed as a heatmap (n = 3).

(K) Intracellular metabolites, as measured by LCMS) of 143B cells expressing empty vector (E.V.) or *Lb*NOX cultured with vehicle or 5 μM AZD7545. Metabolite levels were compared to the E.V. vehicle-treated condition, and the fold change of all metabolites measured were hierarchically clustered and displayed as a heatmap (n = 3).

(L) Principal component analysis of log fold changes in intracellular metabolite levels, relative to vehicle-treated cells, measured in 143B cells cultured with or without 5 μM AZD7545 in the presence or absence of 20 μM duroquinone as indicated (n = 3).

(M) Principal component analysis of log fold changes in intracellular metabolite levels, relative to E.V. vehicle-treated cells, measured in 143B cells expressing empty vector (E.V.) or *Lb*NOX, and treated with vehicle or 5 μM AZD7545 as indicated (n = 3).

(N) Diagram indicating the number of intracellular metabolites altered by AZD7545 treatment in which this alteration is rescued by duroquinone (Duroquinone only), *Lb*NOX expression (LbNOX only), or both (Common metabolites).

Values shown in panels A, C, D, E, F, G, H, and I denote the mean ± SD, and *P* values were calculated by unpaired, two-tailed Student's t-test (n.s. = non-significant).

**Figure supplement 4. PDK inhibition can cooperate with metformin to slow tumor growth.**

(A) Proliferation rate of MDA-MB-361 cells treated with 1 mM metformin, 8 μM AZD7545, and 1 mM pyruvate as indicated (n = 3).

(B) Proliferation rate of HeLa cells cultured in the indicated concentration of metformin and AZD7545, without or with 1 mM pyruvate (Pyr) (n = 3).

(C) Proliferation rate of A549 cells cultured in the indicated concentration of metformin and AZD7545, without or with 1 mM pyruvate (Pyr) (n = 3)

(D) Tumor volume assessment over time for A549 cells implanted in nude mice treated with vehicle, 500 mg/kg metformin, and/or 45 mg/kg AZD7545, administered by daily oral gavage. Treatment was initiated in mice with size-matched tumors upon reaching a volume of 50 $mm^3$. The difference in tumor growth between vehicle and metformin treated mice, and between vehicle and metformin+AZD7545 treated mice was statistically significant (*P* = 0.0014, *P* = 0.0001 respectively by two-way ANOVA). The difference in tumor growth between metformin treated mice and metformin+AZD7545 treated mice approached significance (*P* = 0.0575 by two-way ANOVA) (Vehicle, n = 11; AZD7545, n = 6; Metformin, n = 11; Metformin + AZD7545, n = 8)

(E) AZD7545 concentration in mouse plasma 3 hours (Peak) and 24 hours (Trough) after administration AZD7545 by oral gavage measured by LCMS.

(F) AZD7545 concentration in mouse tumors harvested after three days of AZD7545 treatment by oral gavage as measured by LCMS.

For all panels, values shown denote the mean ± SEM.

**Figure supplement 5. Elevated mitochondrial membrane potential impairs NAD+ regeneration following PDH activation**

(A) Mitochondrial membrane potential, as reflected by TMRE (tetramethylrhodamine, ethyl ester) fluorescence, of primary mouse T cells that were activated by exposure to anti-CD3/CD28 antibodies in the presence of vehicle, 5 μM AZD7545, and/or 500 nM FCCP as indicated. Representative data, as well as the relative median fluorescence values from multiple experiments, are shown (n = 7).

(B) The NAD+/NADH ratio of A549 and HeLa cells following a 5 hour exposure to vehicle or the indicated concentrations of AZD7545 (n = 4).

(C) Relative aspartate levels, normalized to the vehicle-treated condition, of cells treated with vehicle, 2 μM AZD7545, or 2 μM AZD7545 with 250 nM FCCP for 5 hours as measured by LCMS (n = 4).

(D) Proliferation rate of cells cultured with vehicle (V) or AZD7545 in the presence or the absence of 500 nM FCCP (n = 3).

(E) NAD+/NADH ratio of 143B and A549 cells treated with vehicle, 5 μM of AZD7545, or 5 μM of AZD7545 with 0.5 nM gramicidin D (n = 6).

Values shown in all panels denote the mean ± SD, and *P* values were calculated by unpaired, two-tailed Student's t-test. *, **, ***, **** denotes $P < 0.05, 0.01, 0.005, 0.001$; n.s. = non-significant

**Figure supplement 6. Uncoupling NAD+ regeneration from ATP production reduces fermentation in both yeast and mammalian cells.**

(A) Relative pyruvate (Pyr) and lactate (Lac) excretion of 143B cells cultured with or without 20 μM duroquinone normalized to the vehicle-treated condition (n = 3).

(B) Relative proliferation rate of cells cultured in the presence or absence of duroquinone normalized to the vehicle-treated condition. The duroquinone concentration used was 4μM, 16μM, 8μM and 64μM duroquinone for 143B, H1299, C2C12, and PSC cells, respectively (n = 3).

(C) Relative pyruvate (Pyr) and lactate (Lac) excretion of 143B cells expressing empty vector (E.V.) or *Lb*NOX normalized to the E.V. condition. Doxycycline (500 ng/mL) was included in all conditions to induce *Lb*NOX expression (n = 3).

(D) Proliferation rate of primary mouse T cells that had been stimulated with anti-CD3/CD28 antibodies in the presence of vehicle or 250nM FCCP for 1 day (n = 5).

(E) Doubling time of *S. cerevisiae* cultured at different glucose concentrations (n = 3).

(F) Relative ethanol production rate from *S. cerevisiae* assessed at different glucose concentrations. Ethanol production rate is normalized to that of yeast grown in medium containing 0% glucose (n = 3).

(G) Proliferation of *S. cerevisiae* expressing empty vector (E.V.) or *Lb*NOX cultured in standard 3% glucose containing medium (n = 3).

(H) Proliferation of *S. cerevisiae* treated with vehicle or 2 μM FCCP in standard 3% glucose containing medium (n = 3).

Values shown in all panels denote the mean ± SD, and *P* values were calculated by unpaired, two-tailed Student's t-test. n.s. = non-significant
