## Supplemental Figures for "Increased demand for NAD+ relative to ATP drives aerobic glycolysis"

Figure supplement 1

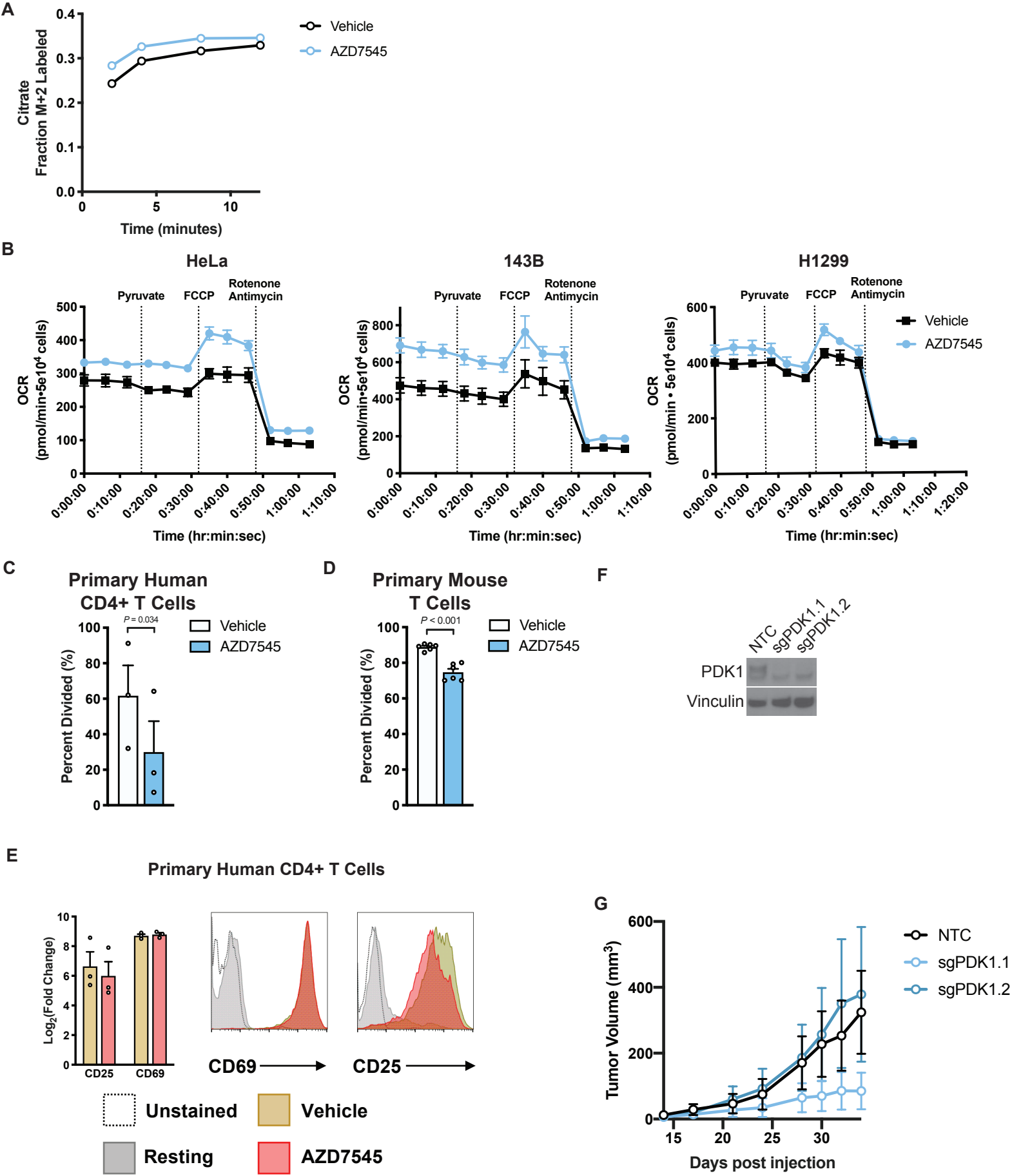

Figure supplement 2

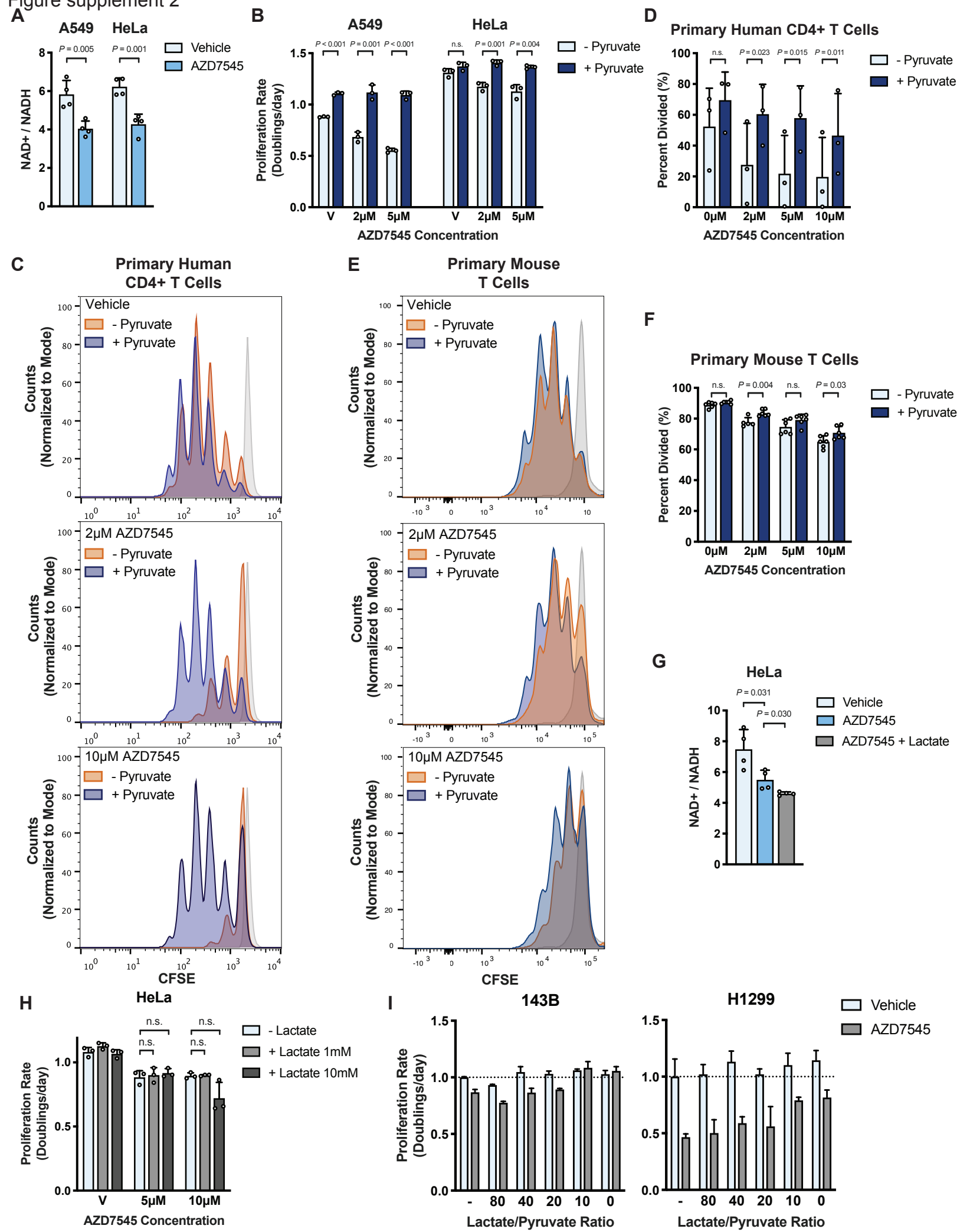

Figure supplement 3

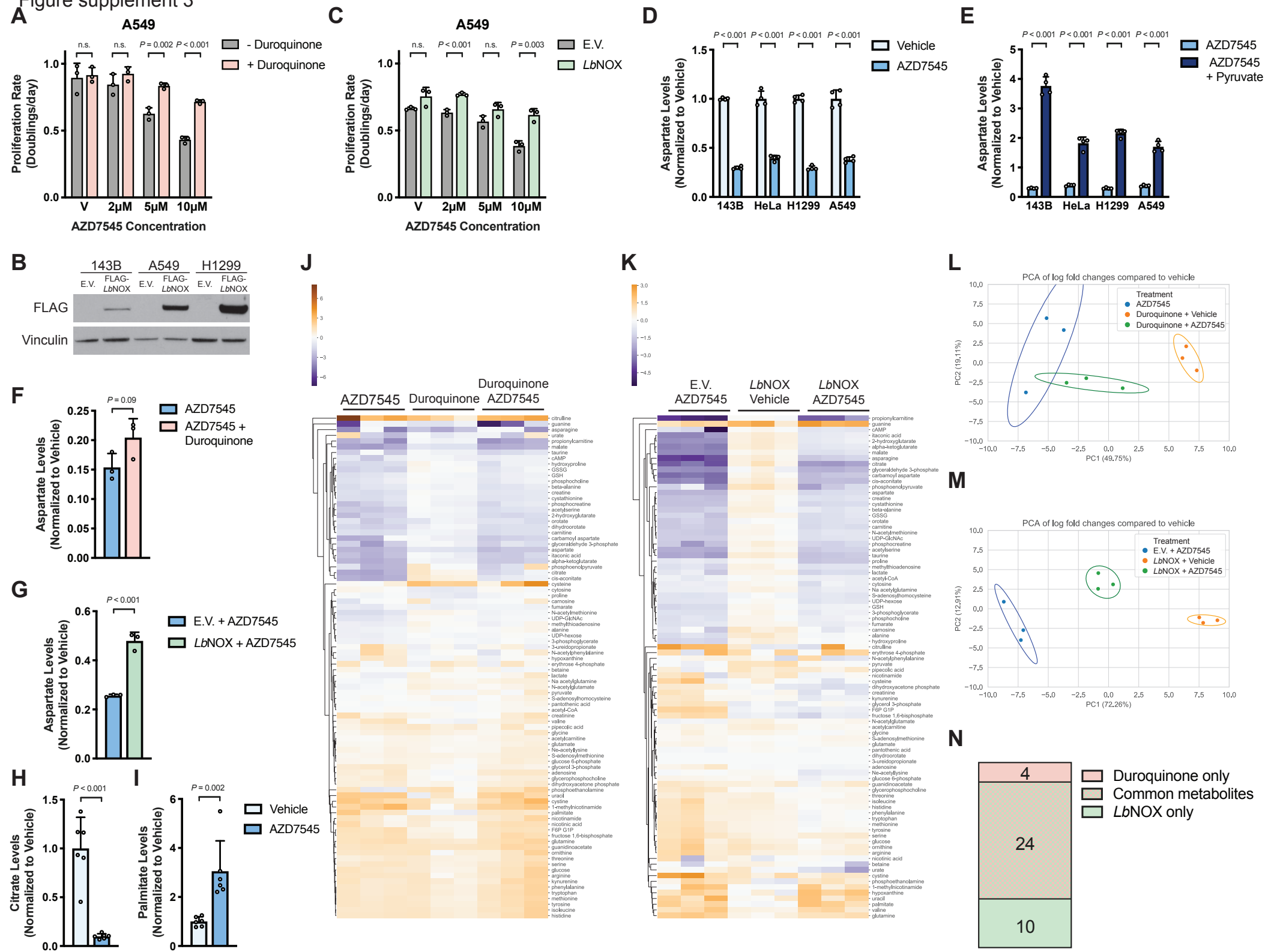

Figure supplement 4

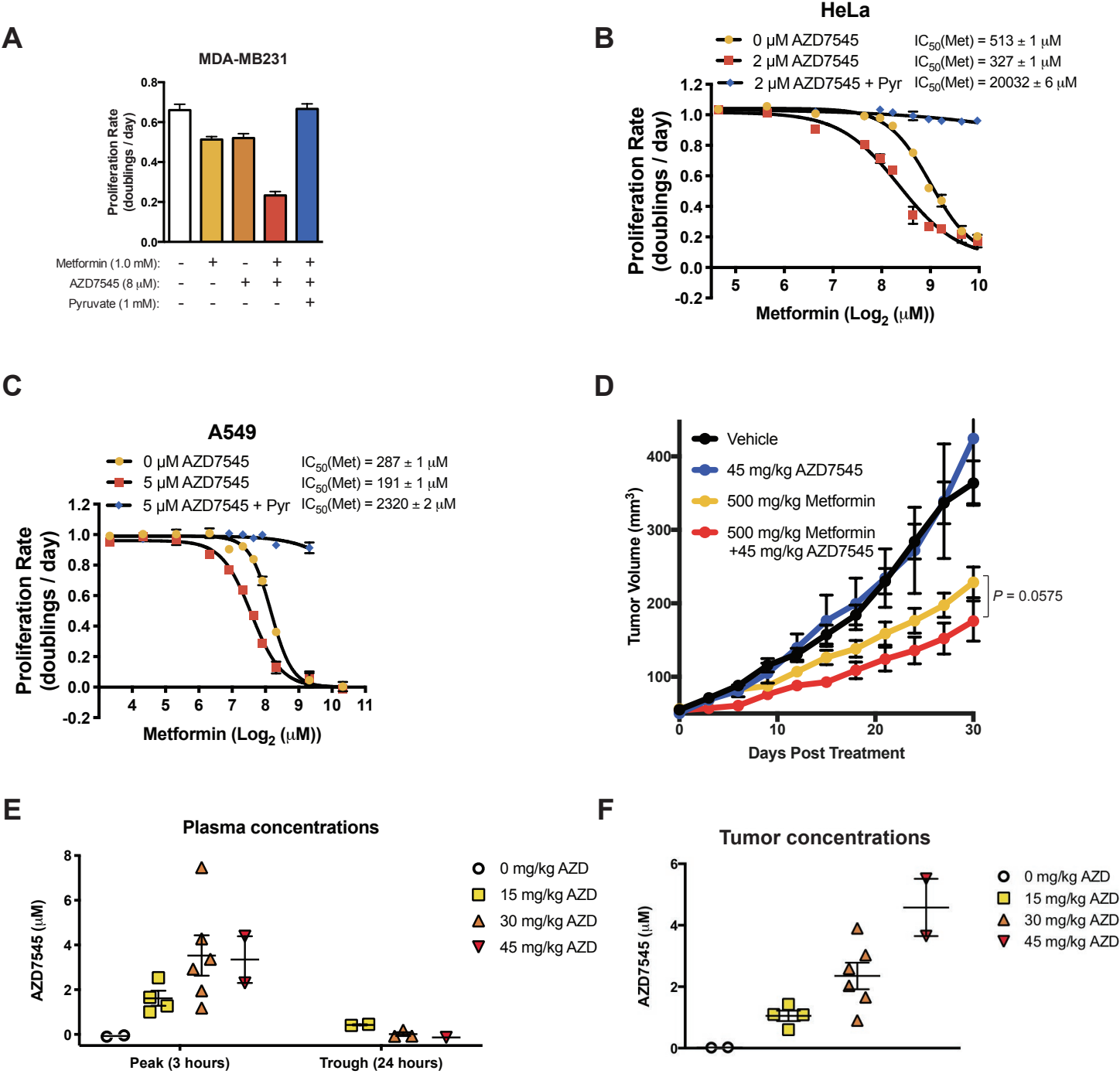

Figure supplement 5

**A**

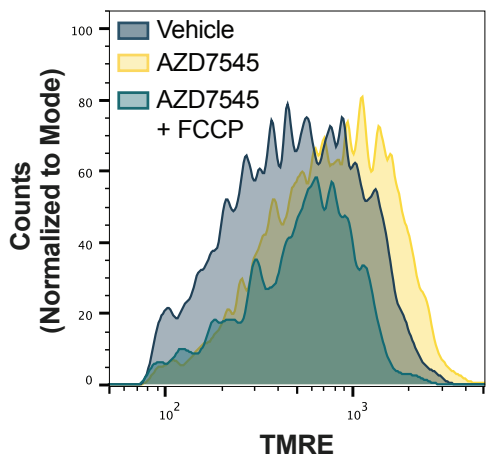

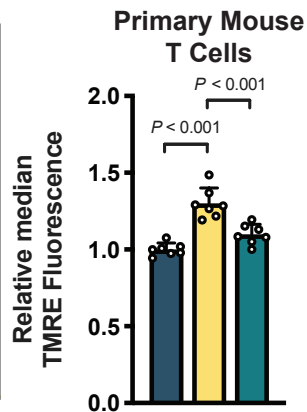

**B**

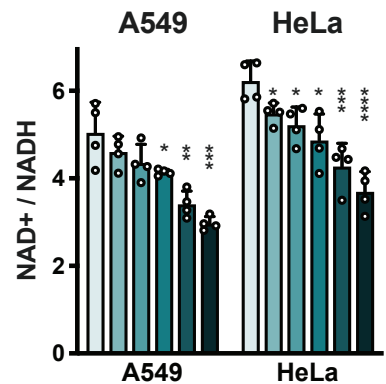

**C**

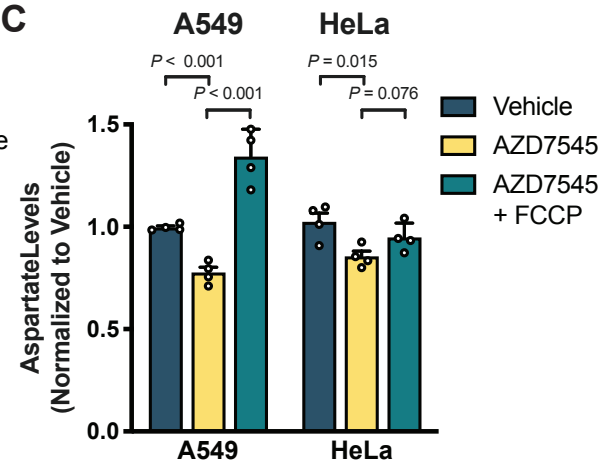

**D**

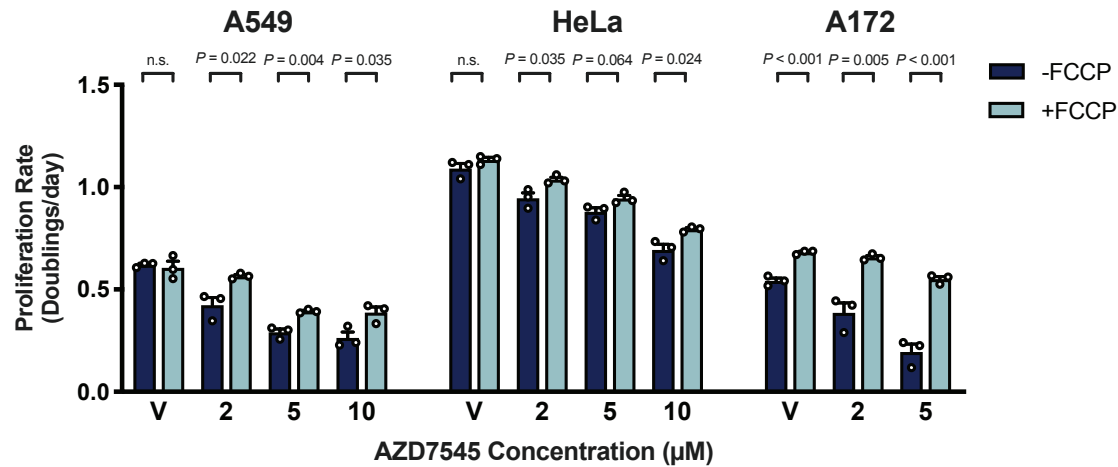

**E**

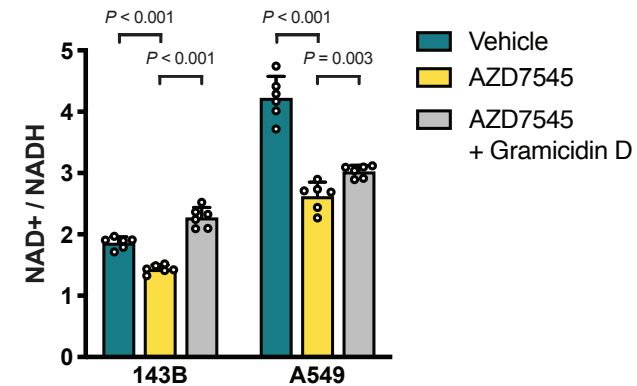

Figure supplement 6

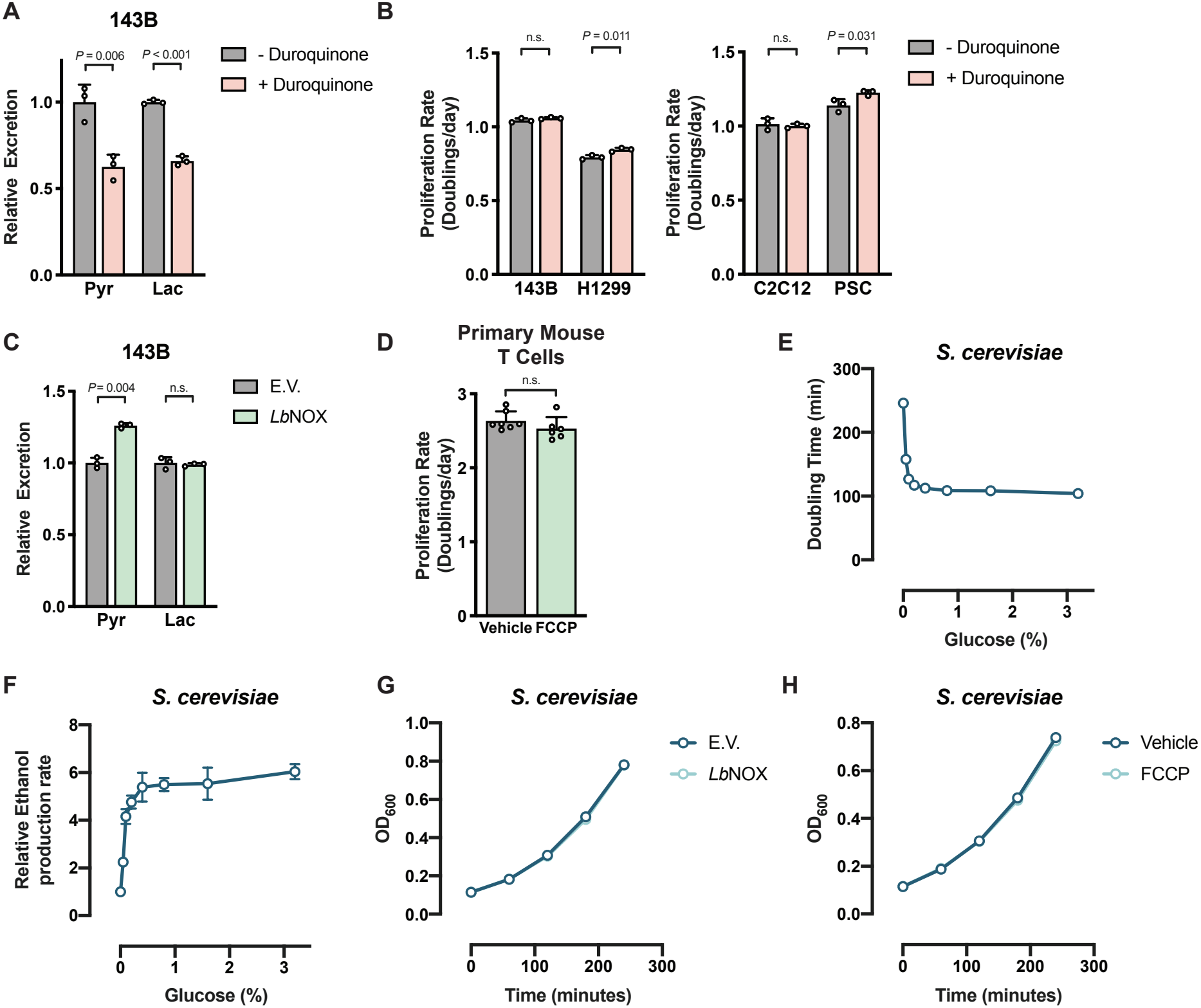
